# Test-Retest Reliability of Dynamic Functional Connectivity Parameters for a Two-State Model

**DOI:** 10.1101/2022.11.15.516555

**Authors:** Xiaojing Fang, Michael Marxen

## Abstract

Dynamic functional connectivity (DFC) metrics derived from resting-state functional magnetic resonance imaging (rs-fMRI) that measure dynamic features of brain communications that change over time are gaining more and more attention. A key concern is the reliability of DFC parameters, especially given the many methodological options. The DFC parameters mean dwell time (*MDT*), prevalence (*Prev*.), inter-transition interval (*ITI*) and state variability (*Var*.) were computed in 23 participants using a two-state model, sliding window analysis, and two imaging sessions. Reliability of the connectivity states, test-retest reliability of the parameters as well as correlations between DFC parameters were investigated for different scan lengths (i.e., 16 vs 8 min.), atlases (i.e., 116 vs. 442 regions of interest) and with/without within-subject centering of DFC matrices. The results showed an integrated (I) and a segregated (S) brain state with high intra-class correlation (ICC) values of the states between sessions (0.67 ≥ ICC ≥ 0.99). Reliability of the DFC parameters is, at best, intermediate with an ICC = 0.51 for prevalence for not-centered, 16 min. data and the coarse atlas. Moreover, *MDT*_s_ of both states were positively correlated for centered data. Prevalence of one state was positively correlated with *MDT* of the same state and frequently negatively correlated with *MDT* of the other state. *Var* was positively correlated with *MDT* and *Prev* for the same state and negatively correlated for the other state for not-centered data. This study suggests that prevalence is the most reliable DFC parameter with lower reliability for shorter scans, more atlas regions and centering. Thus, it is advantageous to formulate hypotheses related to brain dynamics in terms of prevalence, especially in small-scale studies.

## 1. Introduction

The analysis of functional connectivity (FC) between brain regions, a measurement of inter-regional synchrony (Karl J Friston, 1994), has become one of the primary research methods in the field of neuroscience (K. J. Friston, 2011). While many studies implicitly assumed a stationary nature of FC, the dynamic nature of brain activity (Chang & Glover, 2010) is gaining more and more attention in recent years. Dynamic functional connectivity (DFC) describes dynamic features of brain communications that change over time (Hutchison et al., 2013). By now, a large number of works have investigated the dynamic fluctuations of the connections, graphic properties of the DFC matrices and all kinds of characteristics of DFC patterns at rest, illustrating temporal changes in regional interactions, signal transfer between networks and the functional architecture of whole-brain states (Hutchison et al., 2013; Preti, Bolton, & Van De Ville, 2017; Zhang, Baum, Adduru, Biswal, & Michael, 2018). Additionally, a lot of findings revealed not only neural correlates of resting-state DFC with electroencephalography powers (Chang, Liu, Chen, Liu, & Duyn, 2013; Tagliazucchi, von Wegner, Morzelewski, Brodbeck, & Laufs, 2012; Thompson et al., 2013), but also associations of the connectivity with demographics, consciousness, behaviour, cognition (Hutchison et al., 2013; Preti et al., 2017; Shine et al., 2016) and various neurological and psychiatric disorders, such as Parkinson’s disease (Zoller et al., 2019) and schizophrenia (Calhoun, Miller, Pearlson, & Adali, 2014; Hutchison et al., 2013). All the evidence supports that DFC, a reflection of changing patterns of neural activity, is a useful approach to further our understanding of the fundamental characteristics of the brain at rest.

So far, researchers have developed many methods to investigate temporal variations of connectivity, such as time-frequency analysis (Chang & Glover, 2010), time-series modelling (Eavani, Satterthwaite, Gur, Gur, & Davatzikos, 2013; Lindquist, Xu, Nebel, & Caffo, 2014), change-point detecting (Cribben, Wager, & Lindquist, 2013) and co-activation pattern analysis (Liu & Duyn, 2013). Possibly the most popular approach to DFC is sliding window analysis (SWA), which divides a time course into small windows and computes functional connectivity measures for each window (Chang & Glover, 2010). Due to its simplicity in algorithm and application, SWA has been widely used for diverse research interests (Allen et al., 2014; Choe et al., 2017; Hutchison et al., 2013; Preti et al., 2017). However, an inevitable problem of SWA-based DFC measures compared to static FC (SFC) is that reliability is bound to decrease with smaller window size and shorter scan time due to the low signal-to-noise ratio (SNR) of resting-state functional magnetic resonance imaging (rs-fMRI) signals, artefact signals from non-neural factors (e.g., respiration, cardiac pulsation and noise) (Hutchison et al., 2013) and fluctuations of the measures on the time scale of hours and days (e.g. circadian effects) (Orban, Kong, Li, Chee, & Yeo, 2021). As a consequence, sensitivity and specificity of DFC approaches have to be evaluated carefully (Calhoun et al., 2014; Handwerker, Roopchansingh, Gonzalez-Castillo, & Bandettini, 2012; Hlinka & Hadrava, 2015). Hence, an immediate concern is to quantify and maximize the test-retest reliability, measured frequently through intra-class correlation (ICC), of utilized DFC parameters, which will aid to optimize processing pipelines, to register concrete hypotheses, enhance the interpretability of findings and, generally speaking, move this field forward.

Surprisingly, only a few reports of test-retest reliability for brain states were published. By hierarchically clustering the DFC between four subdivisions of the posteromedial cortex and 156 target regions of interest (ROIs), Yang and colleagues observed five clusters in one rs-fMRI session and six in the other for the same 22 participants, revealing partial reproducibility for the states and derived parameters (i.e., duration and transitions) (Z. Yang, Craddock, Margulies, Yan, & Milham, 2014). Another study with 100 subjects from the Human Connectome Project Data Release (HCP) dataset (Van Essen et al., 2013) revealed four similar k-means co-activation clusters across sessions with intermediate reliabilities: ICC ≤ 0.54 for temporal faction (prevalence) or mean dwell time (Smith, Zhao, Keilholz, & Schumacher, 2018). Choe and colleagues (Choe et al., 2017) reported similar values for two k-means connectivity clusters (e.g. ICC = 0.56 for dwell time using SWA). Although there is currently no a priori “correct” number of brain states, when maximizing reliability is the prerogative, choosing k = 2 clusters is advantageous for three reasons: 1) reliability of the state centroid connectivity matrices will be better than for larger k because the states will be most distinct (i.e., furthest away from each other in the clustering space); 2) for parameters such as mean dwell time, noise reduction through averaging should increase with the total time spent in each state (i.e., [scan time]/k, which is largest for k = 2); 3) two states are in line with the the widely accepted view that there are two fundamental principles of brain organization (K. J. Friston, 2009; Shine et al., 2016), namely integration and segregation. This allows a straightforward functional interpretation of the two states.

Naturally, other factors will affect the reliability of DFC parameters such as scan length (Hindriks et al., 2016) and other details of the processing pipelines. Consequently, after validating the interpretation of the two states for our data based on the global graph metrics modularity and global efficiency, we addressed the following questions using a two-state model and a straightforward SWA approach: 1) How reliable are the two states between scanning sessions? 2) How reliable are particular parameters that quantify the dynamics (see section 2.6) for this model? 3) What is the effect of scan length on the reliability? 4) What is the effect of choosing an atlas with a larger number of regions? and 5) What is the effect of within-subject centering of the connectivity matrices before clustering, which will remove differences between subjects in static connectivity?

## 2. Methods

These methods have been described in a similar form for different participants in the preregistration https://osf.io/286fb “Linking Need for Cognition to a Two-State Model of Dynamic Brain Connectivity during Rest”.

### 2.1. Participants

Twenty-five healthy young adults were recruited from a volunteer database. Inclusion criteria were: no magnetic-resonance contraindications, normal or correctable-to-normal vision and an age between 20-35 years old. For the analysis, one participant was excluded because of incomplete MRI data; one participant was excluded because of severe, prefrontal signal drop out despite field map correction. Therefore, we analyzed the data of 23 subjects [mean ± standard deviation (SD) of age: 24.0 ± 3.8 years, 16 females]. Participants received a financial compensation of €40 for full participation in both sessions. The study was approved by the Ethics Committee of the Technische Universtiät Dresden (EK 4012016) and all participants signed informed consent forms after receiving a detailed description of the experiment.

### 2.2. Study Design

To evaluate reliability, a repeated-measure design was used. Study participants were scanned twice with the same scanning protocol with approximately one week between scanning sessions. To eliminate potential session order effects, we generated two sets of data (i.e., session A and B) by randomly assigning the first acquisition session from 12 participants and the second acquisition session from the remaining 11 participants to session A, while the respectively paired scans were assigned to session B.

### 2.3. MRI data acquisition

Imaging data were collected on a Siemens 3 T Magnetom Trio Tim scanner (Siemens, Erlangen, Germany) equipped with a 32-channel head coil. In each session, structural T1-weighted data were acquired after brief localizer scans (~1 min), using a 3D magnetization-prepared rapid gradient echo (MPRAGE) sequence with the parameters of repetition time (TR) 2400 ms, echo time (TE) 2.19 ms, voxel size 0.85 mm × 0.85 mm × 0.85 mm, inversion time 1000 ms, field of view (FOV) 272 mm × 272 mm, flip angle 8°, matrix 320 × 320, slice thickness 1.0 mm, band width (BW) 210 Hz/Px and 240 slices. T2-weighted data were acquired in each session with the parameters of TR 3200 ms, TE 565 ms, voxel size 0.85 mm × 0.85 mm × 0.85 mm, FOV 272 mm × 272 mm, flip angle 120°, matrix 320 × 320, thickness 0.85 mm, BW 744 Hz/Px and 240 slices. We acquired a B0 field map with parameters of TR 832 ms, TE-1 5.19 ms, TE-2 7.65 ms, voxel size 2.0 mm × 2.0 mm × 2.0 mm, gap 0 mm, FOV 208 mm × 208 mm, flip angle 58°, matrix 104 × 104, BW 278 Hz/Px, and 78 slices. Then, we collected rs-fMRI data based on a multi-band axial (T > C ~ −17°) 2D EPI sequence (Moeller et al., 2010) and with the parameters of TR 987 ms, TE 32.6 ms, voxel size 2.0 mm × 2.0 mm × 2.0 mm, slice gap 0 mm, FOV 192 mm ×192 mm, flip angle 62°, matrix 96 × 96, BW 1860 Hz/Px, 72 interleaved slices, and 1000 volumes. The two rs-fMRI scans lasted approximately 16:37 min. each (mean ± SD of the interval between the two scans: 8.87 ± 4.84 days). Additionally, we acquired diffusion-weighted images not relevant to this study. All participants received foam padding for head-movement reduction and earplugs for hearing protection, and were instructed to close their eyes and to try not to fall asleep.

### 2.4. Preprocessing

The following preprocessing pipeline using fMRIPrep 1.2.5 (zenodo.org/record/4252786#.X7TzMGhKhPZ) based on Nipype 1.1.6 (zenodo.org/record/4035081#.X7Ty32hKhPY) (Esteban et al., 2019) was employed: skull stripping, field map correction, co-registration of the structural and functional images, head-motion estimation and correction, spatial normalization to MNI152NLin2009cAsym space. All subjects met our inclusion criteria of less than 7.5% of frames with framewise displacement (Power, Barnes, Snyder, Schlaggar, & Petersen, 2012) ≥ 0.5 mm. Then, the nuisance covariates (i.e., six head motion parameters, signals of cerebral spinal fluid and white matter) were removed from the preprocessed data. The effects of low-frequency drift and high-frequency physiological noise were reduced by using band-pass filtering (1/(*w*TR)* ~ 0.1 Hz) in time (Leonardi & Van De Ville, 2015). Here, we specified *w* (i.e., the window width in TR for the sliding-window analysis) as 40 (Zhang et al., 2018), which resulted in a lower frequency boundary of 0.025 Hz.

### 2.5. Extraction of functional brain states

#### 2.5.1. Sliding-window analysis

In the current study, an instance of a DFC brain state is defined as a connectivity matrix of Fisher-z values based on Pearson’s correlation that were computed for the time window of 40*T*R =* 39.48s based on a previous study (Zhang et al., 2018). The sliding-window analysis was computed in DynamicBC toolbox (Liao et al., 2014) (github.com/guorongwu/DynamicBC) with a step size of 1 TR resulting in 961 brain-state instances (i.e., DFC matrices) per scanning run. Time series were extracted by spatially averaging all voxel-wise signals within each ROI.

#### 2.5.2. Clustering of state instances and back projection

The two-state (i.e., state I and S) classification of the DFC matrices was performed in Matlab 2018b (Mathworks Inc., Natick, MA, USA) by assigning each matrix to one of k = 2 clusters in k-means clustering (Lloyd, 1982) employing cosine distance (*CD)* to measure the similarity between each state instance and the centroids of a cluster (i.e., *CD* = 1 − cos(*α*), where α is the angle between two vectorized DFC matrices. Thus, *CD* ∈ *[0, 2]*, with 0 being the smallest distance, i.e., the highest similarity.) (Karahanoglu & Van De Ville, 2015). Moreover, k = 2 was found to be optimal according to silhouette statistics (Rousseeuw, 1987) computed for cluster numbers k = 2…20 (See Supplemental Fig. S1). For session B, DFC matrices were assigned to state I or S by back-projecting each matrix onto the cluster centroids from session A, i.e. brain state instances from session B were assigned to the state/cluster with the smaller *CD* to the corresponding cluster centroid of session A. Cluster centroids were defined here as the group mean of the normalized, i.e., divided by the root of the sum of squares of all matrix elements, DFC matrices for the respective state in each session. Distances between all state instances and the two cluster centroids were plotted to validate the clustering procedure.

#### 2.5.3. Processing pipeline variations

We investigated 2 × 2 × 2 = 8 different processing pipelines or strategies. Three factors were varied:

1. The length of the rs-fMRI time series: 1000 frames (16:27 min.) versus 500 frames (8:14 min.). For the analysis with 500 frames, we only considered the first 500 frames of a run.
2. The brain atlas used to define the ROIs (see below): 116 ROIs versus 442 ROIs.
3. Within-subject centering of DFC matrices versus ‘not-centered’. For the centered data, the within-subject-and-session mean connectivity matrix was removed from the DFC matrices before the clustering, which eliminates differences in SFC between subjects. As the resulting cluster centroids no longer reflect functional connectivity strength in this case, we added session-level mean connectivity matrices back to the centroid when appropriate.

#### 2.5.4. Employed brain atlases

The Automated Anatomical Labeling atlas (AAL) (Tzourio-Mazoyer et al., 2002) and a joint atlas (Schaefer et al., 2018; Shine et al., 2016) were applied in this study to explore the reliability of dynamic properties of brain states on different spatial scales. The former consists of 116 ROIs and was grouped into nine networks / regions based on Yeo’s seven functional networks of the cerebral cortex (Yeo et al., 2011) and anatomical parcellations of the subcortical structures and the cerebellum: the visual network, the sensory-motor network, the dorsal attention network, the ventral attention network, the limbic network, the fronto-parietal network, the default mode network, the basal ganglia network, and the cerebellar network. The joint atlas consisted of 442 ROIs: 400 cortical parcels of the Schaefer atlas (Schaefer et al., 2018), 14 subcortical regions from the Harvard-Oxford subcortical atlas, i.e. the bilateral thalamus, caudate, putamen, <u>ventral striatum,</u> globus pallidus, amygdala, and hippocampus, and 28 cerebellar regions from the SUIT atlas (Diedrichsen, Balsters, Flavell, Cussans, & Ramnani, 2009). This joint atlas was grouped into the same nine networks used for the AAL atlas (see Supplementary Tables S1-S3).

### 2.6. DFC parameters

We computed seven different dynamical parameters of the brain based on the state sequence of each subject in each session (i.e., all the dynamic parameters were specific to each of the two states and were indexed by I or S, except *ITI*):

a. Mean dwell time (*MDT*): the mean time one participant maintained state I or S continuously during a run. Note that we excluded all windows within the very first and last states from the computation of *MDT* since the beginning/end of the state is unknown. Consequently, only individual DFC series with at least two state transitions were used to calculate *MDT* for at least one of the states.
b. Prevalence (*Prev*.): the fraction of windows spent in state I or S concerning with respect to the total number of windows in the DFC series. Note: *Prev*_*I*_ = 1 − *Prev*_*S*_.
c. Inter-transition interval (*ITI*): the average time length remaining in any state before transitioning to the other state during scans, i.e., *ITI* = (*MDT*_*I*_ + *MDT*_*S*_)/2 for an even number of states in a series. Note that only DFC series including at least three state transitions were considered for the computation.
d. State variability (*Var*.): the mean Euclidean distance of each state instance of one subject to the run-average of all state instances within the same state:

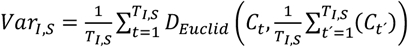, where *C*_*t*_ is the DFC matrix at time window *t* and *D*_*Euclid*_(*C*_*a*_, *C*_*b*_) the Euclidean distance between the vectorized DFC matrices *C*_*a*_ and *C*_*b*_. Note that this definition does not use cosine distance or the above definition of the cluster centroid because such metrics are not invariant to translations in the clustering space.

Following the above definitions, the seven parameters are not completely independent. Nevertheless, each of them represents a different aspect of brain state dynamics and may maximize associations with other parameters. The number of subjects used to compute each parameter are given in supplementary table S5.

### 2.7 Data analyses

#### 2.7.1 Reliability of cluster centroids

To evaluate the reliability of cluster centroids between sessions, we computed the intra-class correlation coefficient (ICC) (Bartko, 1966), a popular measurement that quantifies test-retest reliability and was previously used in other rs-fMRI studies (Choe et al., 2017; Zhang et al., 2018; Zuo & Xing, 2014):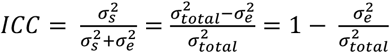, where, 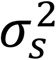 indicates the between-subject variance, 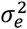 the within-subject variance and 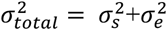 the total variance observed in the data.

The ICC used in this study was based on a one-way ANOVA intra-class correlation framework (Bartko, 1966). Note that ICC values may become less than zero in this framework if variability within the group is greater than across groups (i.e., “divergence” within a group). In addition, we calculated Spearman’s correlation, which is robust to outliers and non-normal distributions (de Winter, Gosling, & Potter, 2016) and would quantify reliability of within-session ranks.

#### 2.7.2 Graph-theoretical analyses of cluster centroids

To describe the functional organization characteristics of the two brain states (i.e. cluster centroids), we computed the two global graph parameters modularity and global efficiency (Cohen & D’Esposito, 2016; Rubinov & Sporns, 2010) for the 16-min data in brain connectivity toolbox (BCT, Version 2019-03-03, nitrc.org/frs/?group_id=241):

a. Modularity 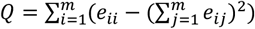, where *e_ij_* means the proportion of edges that link a node in module *i* to any node in module j, and *m* the number of all non-overlapping modules generated by Newman’s modularity algorithm (Newman & Girvan, 2004).
b. Global efficiency 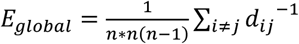, where n is the number of nodes, and d_ij_ the minimum distance between node i and j. In this study, we only computed the distance based on connectivity between the nodes with positive strength (Barch et al., 2013).

We evaluated the variance within these graph measures by generating 1000 bootstrapping samples of the DFC matrices for each state/cluster of session A.

#### 2.7.3 Reliability of dynamic parameters

We computed values of ICC and Spearman’s correlation to quantify the reliability of the dynamic parameters between the two sessions. Confidence intervals of the reliability were estimated by bootstrapping with 1000 samples.

#### 2.7.4 Correlations between DFC parameters

To quantify the interdependence between the dynamic parameters, we calculated Spearman’s correlations between the averaged parameter values across the two sessions.

## 3. Results

### 3.1. Reliability of cluster centroids

Results of the clustering procedure (session A) and back-projection (session B) for all processing strategies are shown in Fig. 1 for the coarse atlas and Fig. S2 for the fine atlas, consistently demonstrating that the DFC matrices cluster around an overall strongly connected (i.e., integrated) state I and a weakly connected (i.e., segregated) state S (see also Section 3.2). For the centered data (Fig. 1), the centroid of state I shows correlations above the run-mean while state S shows correlations below the run-mean (see also Fig. S5). Quantitative reliability between the clusters from session A and B in terms of *ICC* and Spearman’s correlation are given in Table 1 and Table S6, respectively. Note that, for the not-centered data, reliabilities were also computed after removing the group-mean matrix (Table 1c and Table S6c) to eliminate the bias that not-centered centroids generally exhibit high positive ICC values even between different states (Table 1a and Table S6a). Overall, the reliabilities between corresponding states were generally high (ICC ≥ 0.67), especially for longer scan time (ICC ≥ 0.86), while not-corresponding states showed negative ICC values in the centered data. Cosine distances between corresponding states were generally very low (CD ≤ 0.08) and very high for not-corresponding states (CD ≥ 1.96) in the centered data (see Table S6). State reliability generally decreased for shorter scan lengths, more ROIs and centering.

**Table 1.** Reliability (measured by ICC) between the two clusters for the two sessions

| 116-ROI atlas |  |  |  |  |  | 442-ROI atlas |  |  |  |
| --- | --- | --- | --- | --- | --- | --- | --- | --- | --- |
| a) Clusters based on not-centered data |  |  |  |  |  |  |  |  |  |
|  |  | 16-min data |  | 8-min data |  | 16-min data |  | 8-min data |  |
|  |  | Session B |  | Session B |  | Session B |  | Session B |  |
|  |  | State S | State I | State S | State I | State S | State I | State S | State I |
| Session A | State S | ICC = 0.97 | ICC =0.71 | ICC = 0.96 | ICC =0.72 | ICC = 0.96 | ICC =0.57 | ICC = 0.94 | ICC =0.61 |
|  | State I | ICC = 0.67 | ICC =0.99 | ICC = 0.66 | ICC =0.98 | ICC = 0.55 | ICC =0.98 | ICC = 0.55 | ICC =0.96 |
| b) Clusters based on centered data |  |  |  |  |  |  |  |  |  |
|  |  | 16-min data |  | 8-min data |  | 16-min data |  | 8-min data |  |
|  |  | Session B |  | Session B |  | Session B |  | Session B |  |
|  |  | State S | State I | State S | State I | State S | State I | State S | State I |
| Session A | State S | ICC = 0.93 | ICC =-0.14 | ICC = 0.86 | ICC =-0.13 | ICC = 0.86 | ICC =-0.07 | ICC = 0.73 | ICC =-0.07 |
|  | State I | ICC = -0.13 | ICC =0.94 | ICC = -0.12 | ICC =0.84 | ICC = -0.06 | ICC =0.87 | ICC = -0.06 | ICC =0.67 |
| c) Clusters based on not-centered data & removing session mean matrices |  |  |  |  |  |  |  |  |  |
|  |  | 16 min data |  | 8 min data |  | 16 min data |  | 8 min data |  |
|  |  | Session B |  | Session B |  | Session B |  | Session B |  |
|  |  | State S | State I | State S | State I | State S | State I | State S | State I |
| Session A | State S | ICC = 0.93 | ICC =-0.47 | ICC = 0.86 | ICC =-0.32 | ICC = 0.89 | ICC =-0.22 | ICC = 0.79 | ICC =-0.19 |
|  | State I | ICC = -0.47 | ICC =0.93 | ICC = -0.32 | ICC =0.86 | ICC = -0.22 | ICC =0.89 | ICC = -0.19 | ICC =0.79 |

**Fig. 1.**
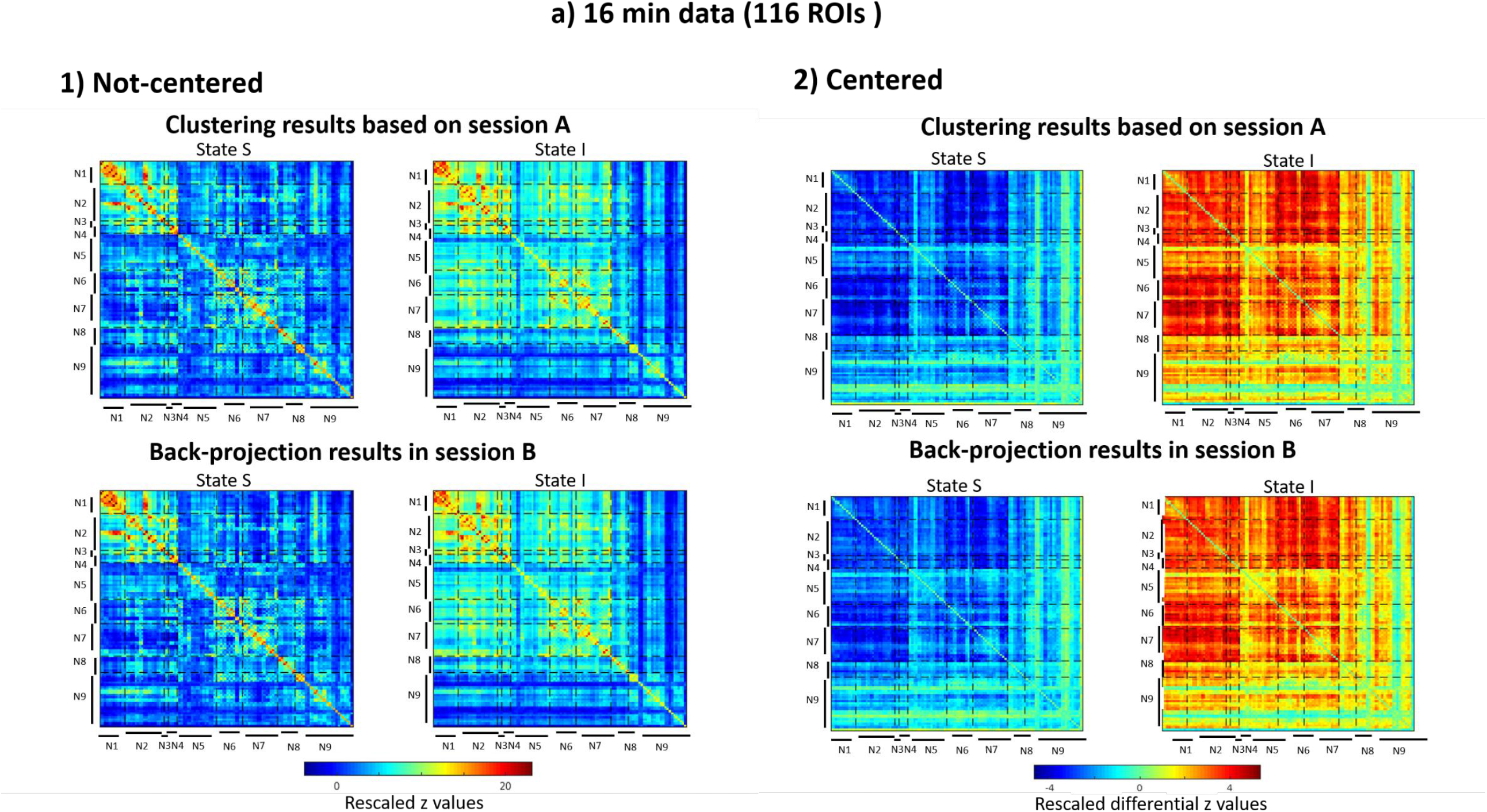
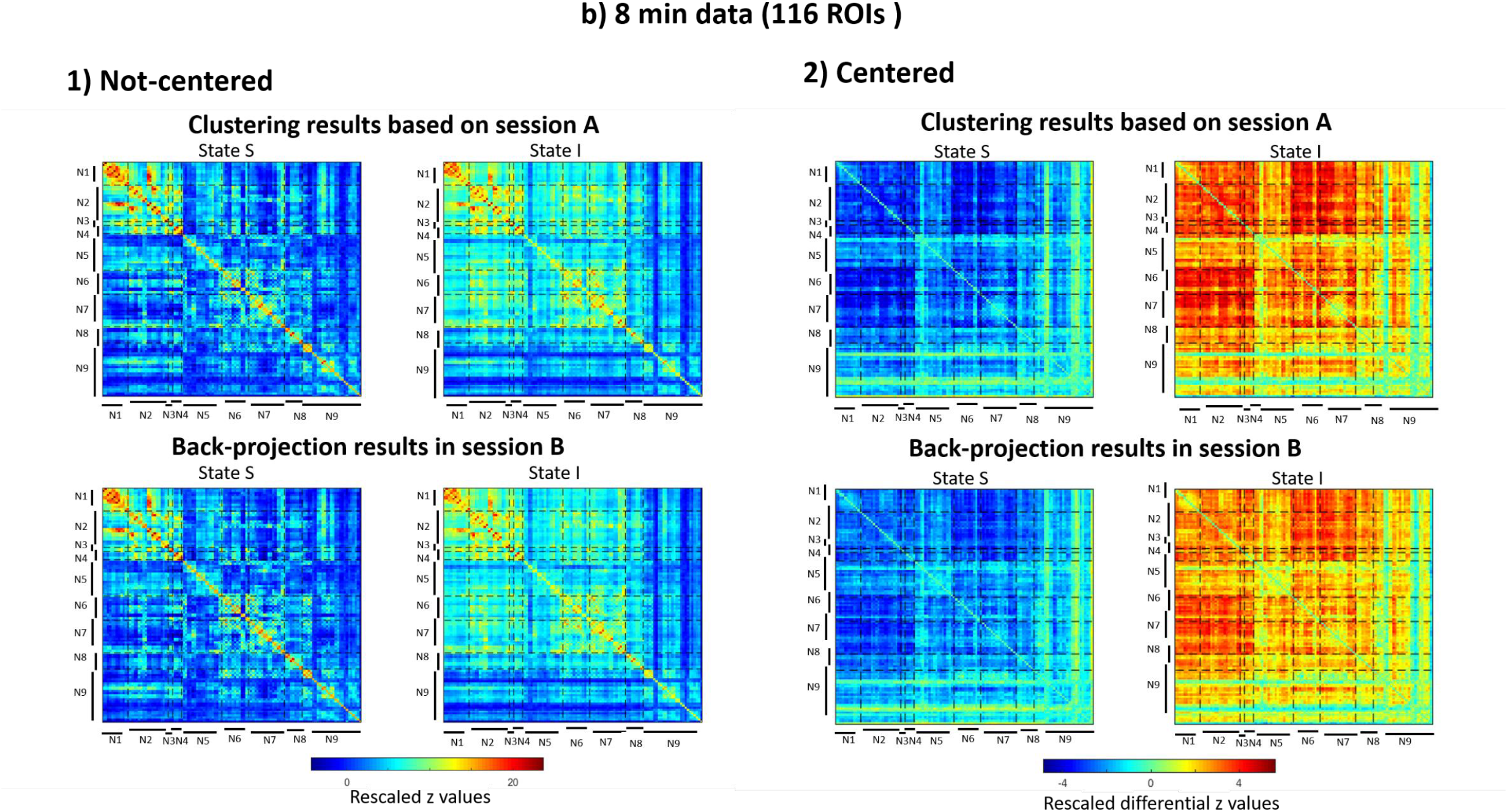
Cluster centroids (k = 2) of the DFC matrices based on the AAL atlas for different time-length and not-centered/centered data. For session A, k-means clustering was used for the state assignment, while back projection results are shown for session B. Displayed are the cluster centroids (see Methods; N1: visual network; N2: sensory-motor network; N3: dorsal attention network; N4: ventral attention network; N5: limbic network; N6: fronto-parietal network; N7: default mode network; N8: basal ganglia network; N9: cerebellum network).

**Fig. 2.**
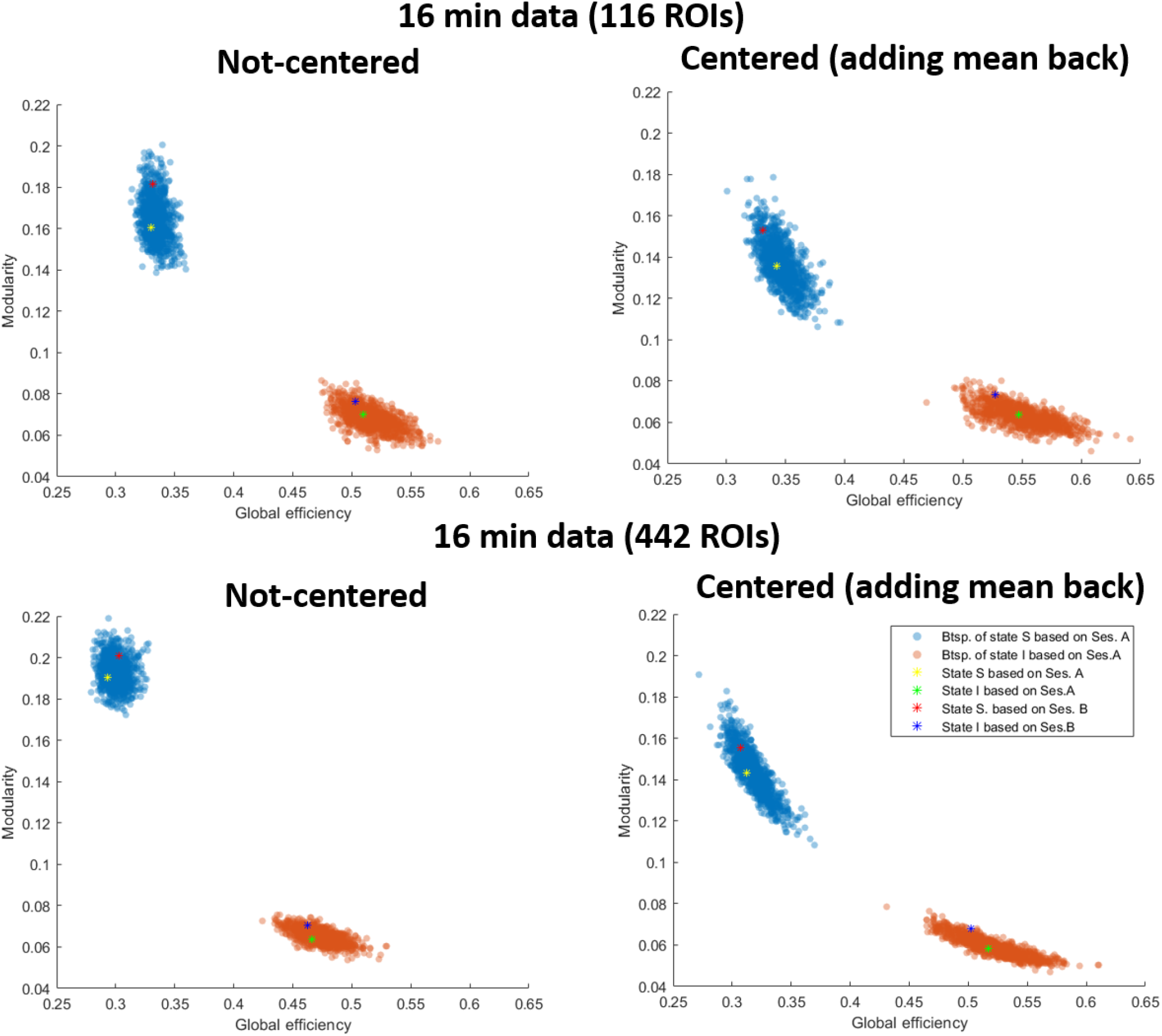
Global graph-theoretical measures of cluster centroids for 16-min data. Point clouds were generated by bootstrapping DFC matrices from session A and recalculation of the cluster centroid (Btsp. - bootsptraping; Ses. – session). For centered data, the session mean was added back. Note that session B centroids were computed as mean values of the respective state after back projection (see Methods).

### 3.2. Graph-theoretical Analyses

The two global graph parameters modularity and efficiency are shown in Fig. 5 for the 16-min data including results of the bootstrapping procedure. The results demonstrate that the graphs of the two states were distinct with respect to these two parameters: the state with low connectivity, i.e., the “segregated” state (S), had low global efficiency and high modularity, while the state with strong connectivity, i.e., the “integrated” (I) state, had high global efficiency and low modularity. Bootstrapping for session A showed that these clusters did not overlap within this two-parameter space. Additionally, the modularity and global efficiency values of the cluster centroids in session B were in the range of the bootstrapping results for session A (Fig. 5; Table S4). Note that individual centering reduced modularity and increased global efficiency for both states (Table S4). Additionally, the two clouds moved closer to each other for the centered data (Fig. 5).

### 3.3. Reliability of dynamic parameters

ICCs, *p*-values and confidence intervals based on bootstrapping for all parameters and pipelines are given in Table 2. Spearman’s correlation are given in Table S7. The highest ICC value of 0.55 was obtained for *Var*_*I*_ for the 8-min data, however, it appears to be driven by outliers as the Spearman correlation is far from significant and the confidence interval is very wide. The next highest ICC value of 0.51 was obtained for prevalence for the 16 min. data with the coarse atlas and without centering (Fig. 3). It is the only parameter that shows consistent values between pipelines at an intermediate level (0.22 ≤ ICC ≤ 0.51) with a number of significant values (uncorrected).

**Table 2.**
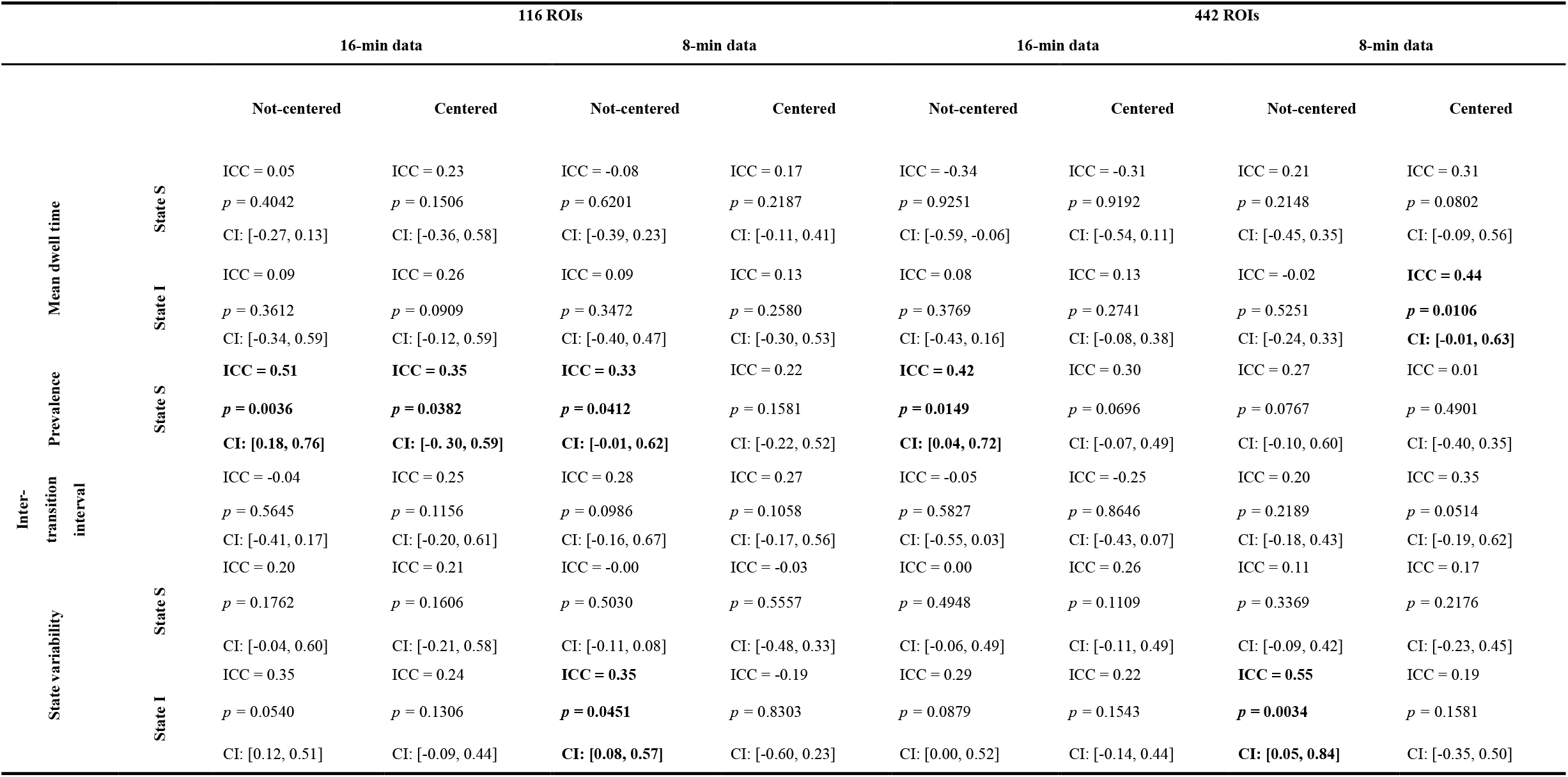
ICC values of the dynamic parameters for all pipelines. CI: confidence interval based on bootstrapping (1000 times). Bold: significant reliability (uncorrected).

**Fig. 3.**
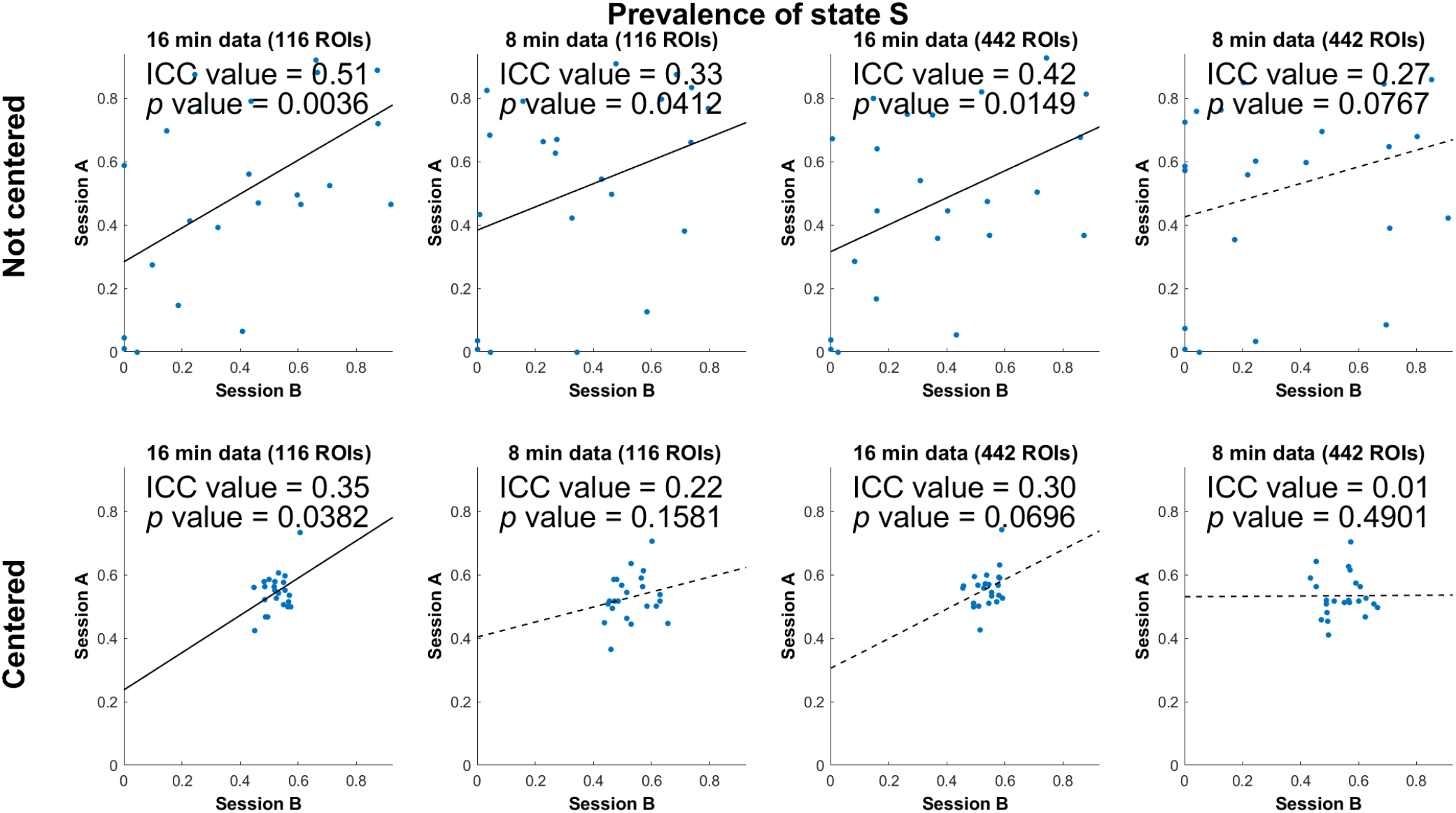
Scatter plots of prevalence between sessions for different pipelines to illustrate reliability. Solid line: *p* value < 0.05; dotted line: *p* value > 0.05.

### 3.4. Correlations between DFC parameters

Spearman’s correlations between different dynamic parameters are given in Tables 3 and 4 for the coarse atlas and tables S8 and S9 for the fine atlas. With some consistency over all pipelines (but not always significant), *MDT*_s_ of both states were positively correlated for centered data. Prevalence of one state was positively correlated with *MDT* of the same state and frequently negatively correlated with *MDT* of the other state. *Var* was positively correlated with *MDT* and *Prev* of the same state and negatively correlated with *MDT* and *Prev* of the other state for not-centered data. As to be expected, *ITI* was positively associated with the *MDT*_s_, especially with *MD_s_* and in the 16-min data.

**Table 3.**
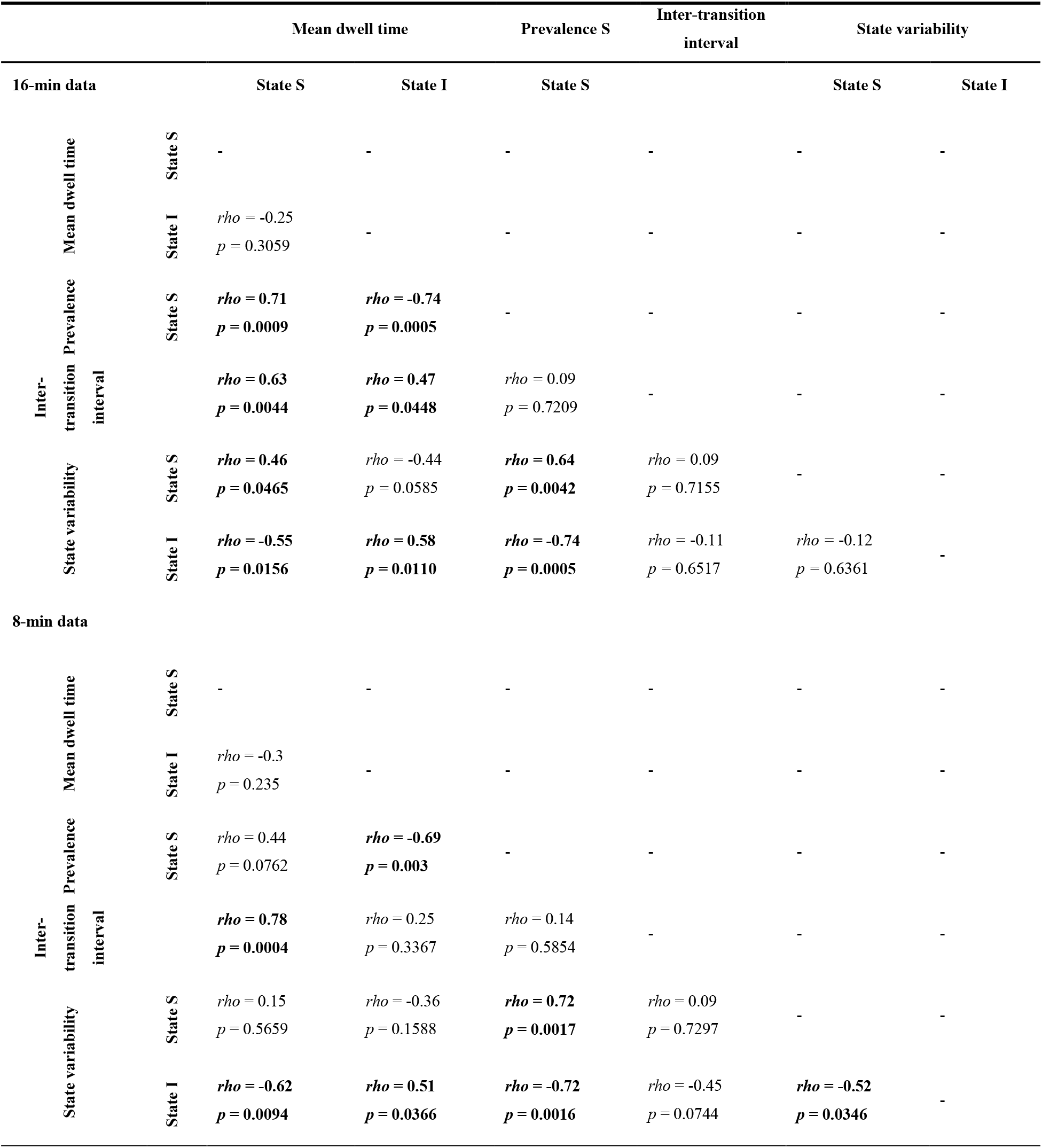
Spearman’s correlations between DFC parameters (mean of both sessions) for not-centered data and 116 ROIs (AAL atlas). Bold: significant correlation (uncorrected).

**Table 4.**
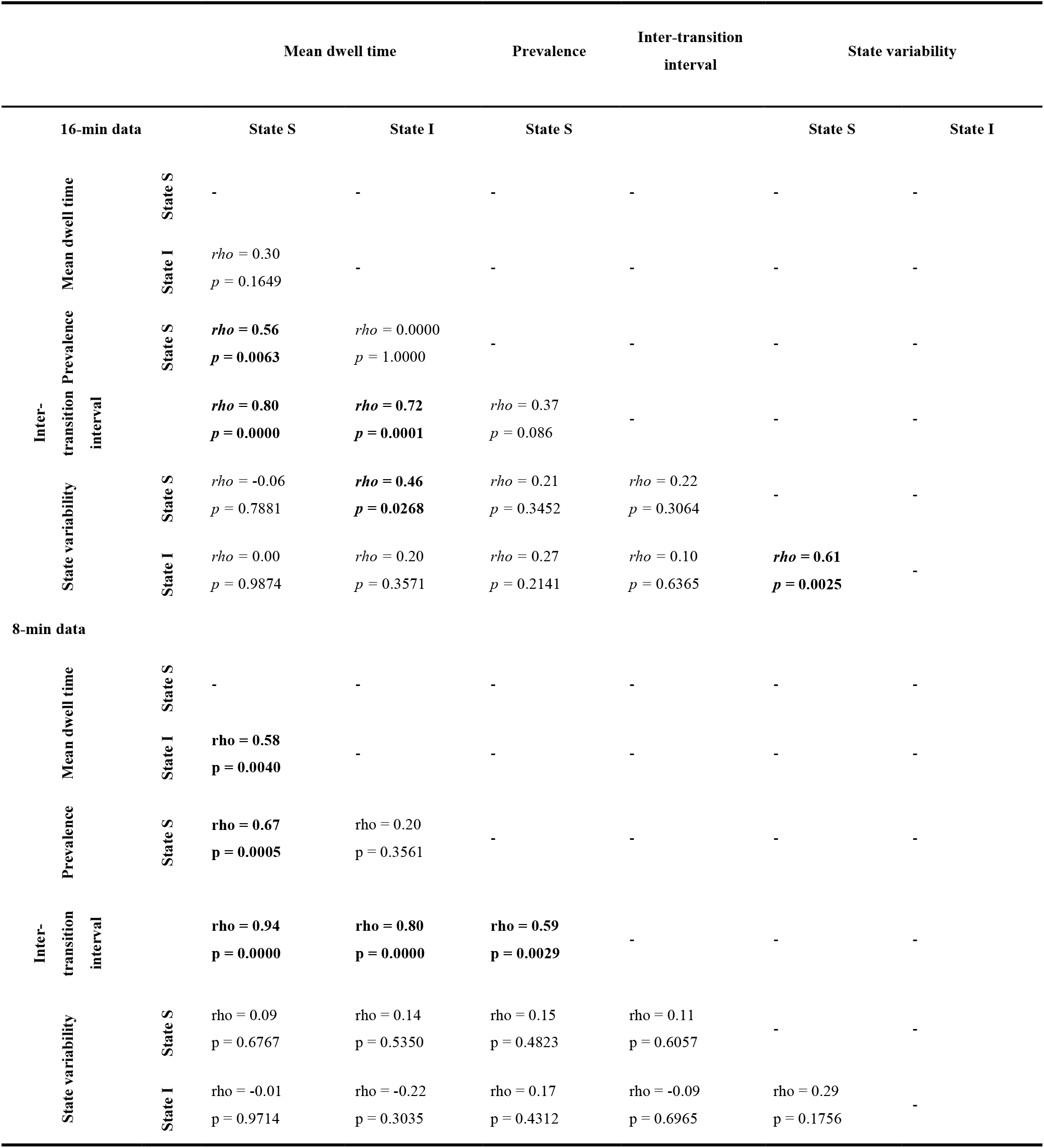
Spearman’s correlations between DFC parameters (mean of both sessions) for centered data and 116 ROIs (AAL atlas). Bold: significant correlation (uncorrected).

## 4. Discussion

To evaluate the potential utility of DFC parameters as personality traits or biomarkers for further correlation analyses, knowing and maximizing their reliability can be crucial. In the introduction, we made an argument for a two-state model as a good starting point. Our graph-based analysis confirmed our expectation that clustering results in an integrated brain state with low modularity and high global efficiency and a segregated brain state with opposite properties. Subsequently, we investigated the reliability of the resulting state centroids and seven parameters using eight different pipelines. We used rs-fMRI data of 23 subjects from two sessions using an in-house fMRI sequence protocol with multi-band factor 6, an isotropic resolution of 2 mm and a TR = 0.987s. This sequence was also used in other studies of our center, which is one reason for us to evaluate reliability of parameters derived from this particular sequence. Because of the relatively small sample size, some of our observations need to be regarded with care, as the differences between pipelines, for example, are not statistically significant.

### Reliability of cluster centroids

Consistent with Choe and colleagues’ study (Choe et al., 2017), we found high reliability of the paired clusters between sessions for both states based on all the strategies. However, shorter scan times and a finer atlas both reduced the ICC reliability from 0.93 to 0.67 (Table 1) for the centered data. This effect can also be observed when looking at the similarity of the clusters in terms of cosine distance (Table S6). For the not-centered data after removing the session mean, the effect was not as severe dropping ICC from 0.93 to 0.79. These effect needs confirmation in a larger study. As to be expected, removing the session mean from the not-centered data increased cosine distances of the unpaired clusters, because the vectors are moved towards the center of the clustering space.

Furthermore, the structure of the two states based on the not-centered data is visually consistent with dynamic two-state patterns in the whole brain at rest for not only healthy people (Choe et al., 2017) with eyes opened/closed (Weng et al., 2020) and different ages (young as 20-35 years old and older as 55-80 years old)(X. Yang et al., 2022), but also clinic patients of Parkinson’s disease (Fiorenzato et al., 2019; Kim et al., 2017), major depressive disorder (Zheng et al., 2022) and dementia with Lewy bodies (Ma et al., 2019). This consistency in state structure between studies is encouraging because, if the structure had varied, statements about brain organization would have lacked a common ground for comparison. However, this comparability may be severely reduced for short scan lengths and fine atlases. Also, note that, for the not-centered data without removing the session mean, ICC values overestimate reliability in the sense that ICC values are even high between the two different states (ICC ≤ 0.72), which indicates that the two states differ primarily in strength, not the direction of the correlations. Figures S3 and S6 show the effect of centering on the distances to each centroid. Centering increases the dissimilarity between the unpaired states. It also shifts the prevalence of either state towards 0.5 (see Fig. 3) and, thus, reduces the number of subjects that remain in the same state for the whole scanning run, i.e., never change states.

### Reliability of dynamic parameters

*Prev* showed the highest reliability across analyses (i.e., 4/8 pipelines), which is consistent with existing findings (Abrol et al., 2017; Choe et al., 2017; Smith et al., 2018) and our theoretical consideration that it is computed based on the complete scan time, while, for example, *MDT* is derived in average from only half the scan time. Thus, the *Prev* parameter is recommended for the formulation of hypotheses related to correlations between dynamic measures of functional brain integration and other subject characteristics.

Reliability of the DFC parameters is, at best, intermediate with an ICC = 0.51 for prevalence for not-centered, 16 min. data and the coarse atlas. Shorter scans, more atlas regions, and centering reduced this value. The reliability of the *MDT*_s_ was less than of prevalence with a maximum of ICC = 0.44 for the centered, 8 min. data with the fine atlas, which was expected for the above reason. The finding that centering increased the reliability of *MDT* for all pipelines indicates that between-session variations in SFC have a negative impact on the reliability of *MDT*_s_, which seems to be reversed for prevalence where between-subject variations in static connectivity (Zurich, 2017) appear to help with re-identification of a subject in the second session.

Several reports suggested a higher reproducibility (Abrol et al., 2017) and higher reliabilities (ICC = 0.56 (Choe et al., 2017) and ICC ≤ 0.47 (Smith et al., 2018)) of the *MDT*_s_ than observed by us. This inconsistency suggests a high sensitivity of *MDT* to differences in methodological (e.g., direct clustering versus two-level clustering and ROIs selection based on independent component analysis versus parcellation atlas) (Abrol et al., 2017; Choe et al., 2017; Smith et al., 2018). In addition to the MRI sequence parameters, both the number of regions as well as the ICA-based noise filtering may have an impact here. The reliabilities of the *ITI* and *Var* parameters were low and not significant for many pipelines and not systematically related to the pipelines. More power would be needed to make further claims about these parameters.

Several considerations are relevant with respect to centering. To obtain the highest reliability for prevalence, centering is not recommended. It will eliminate a large portion of the between-subject variance, which is a disadvantage for cross-sectional analyses. It will also effect the interpretation of the brain state and may make the labels “integrated” versus “segregated” less meaningful, i.e., a subject that displays more integration (high connectivity) over all frames may lose this label when such between-subject differences in overall (static) connectivity are removed. Not to center, however, also has considerable disadvantages: 1) as seen, it reduces reliability of the *MDT*_s_; 2) without centering, static and dynamic effects are mixed, i.e. between-subject differences may be driven by differences in static connectivity rather than by the dynamics; and 3) centering could reduce noise if differences in static connectivity are fluctuating on the time scale of hours to days and do not effect the dynamics. Thus centering may be especially recommended for longitudinal studies over longer time periods.

### Correlations between DFC parameters

*MDTs* of both states were positively correlated for centered data between the two states and with *ITI*, which indicates a tendency for subjects towards less state switching with longer *MDT*_s_ rather than a compensatory or anti-correlated pattern that a longer duration of *MDT*_*s*_ would be associated with shorter *MDT*_*I*_. The result revealed that prevalence of one state was positively correlated with *MDT* of the same state and frequently negatively correlated with *MDT* of the other state. This indicates that statements relating to brain integration may be either put in terms of prevalence or *MDT*_*I*_, i.e. a more integrated brain would have both a higher prevalence and longer*MDT*_*I*_. *Var* was positively correlated with *MDT* and *Prev* for the same state and negatively correlated for the other state for non-centered data. However, we need to carefully interpret these results given the low reliability of the *Var* parameter.

### Limitations and future works

There are some limitations of the current work that should be addressed in the future: 1) as mentioned above, the primary limitation of the current study is the small sample size, which does not allow us to observe significant differences between pipelines and parameters. Thus, we are preparing to repeat the analysis in a larger cohort of Human Connectome Project (www.humanconnectome.org) subjects. 2) We employed back-projection in the second session instead of re-clustering to avoid mixing effects of clustering due to noise on the group level and differences in connectivity. It is unclear whether re-clustering would introduce noise or remove it. To answer this question, one would need to repeat the analysis with re-clustering. 3) The two-state model may not be optimal to maximize reliability of a specific state or identify group differences in such specific states. Thus, models with more states and other dynamic methods, e.g. co-activation analyses (Karahanoglu & Van De Ville, 2015), could show superiority in detecting specific differences in different networks. 4) Our clustering approach does not allow for mixed states. Several studies suggested that a subject may simultaneously be in multiple overlapping states (Leonardi, Shirer, Greicius, & Van De Ville, 2014; Miller et al., 2016). 5) Many more processing pipelines are imaginable that may improve reliability, e.g. more atlases with different granularities, different noise filters, different MR-sequences, co-activation versus temporal correlation.

## Conclusions

In summary, we found high reliability of clustering results with a two-state model but only medium to low reliability of the dynamic parameters. We recommend state prevalence as the most reliable parameter to formulate hypotheses. Shorter scan lengths and a finer atlas decrease reliability while centering reduces reliability of prevalence but improves reliability of *MDT*_s_.

Larger samples are needed to substantiate our findings.

## Funding statement and acknowledgements

This research was supported by the Deutsche Forschungsgemeinschaft (DFG) grants 178833530 (SFB 940) and 402170461 (TRR 265).

