## Supplementary tables and figures for "Test-Retest Reliability of Dynamic Functional Connectivity Parameters for a Two-State Model"

**Table S1.** Functional network mapping of AAL parcels into nine networks

| ROI name | ROI Label | Network |
|---|---|---|
| Calcarine_L | 43 | Visual network |
| Calcarine_R | 44 | Visual network |
| Cuneus_L | 45 | Visual network |
| Cuneus_R | 46 | Visual network |
| Lingual_L | 47 | Visual network |
| Lingual_R | 48 | Visual network |
| Occipital_Sup_L | 49 | Visual network |
| Occipital_Sup_R | 50 | Visual network |
| Occipital_Inf_L | 53 | Visual network |
| Occipital_Inf_R | 54 | Visual network |
| Fusiform_L | 55 | Visual network |
| Fusiform_R | 56 | Visual network |
| Precentral_L | 1 | SM network |
| Precentral_R | 2 | SM network |
| Rolandic_Oper_L | 17 | SM network |
| Rolandic_Oper_R | 18 | SM network |
| Supp_Motor_Area_L | 19 | SM network |
| Supp_Motor_Area_R | 20 | SM network |
| Cingulum_Mid_L | 33 | SM network |
| Cingulum_Mid_R | 34 | SM network |
| Occipital_Mid_L | 51 | SM network |
| Occipital_Mid_R | 52 | SM network |
| Postcentral_L | 57 | SM network |
| Postcentral_R | 58 | SM network |
| Paracentral_Lobule_L | 69 | SM network |
| Paracentral_Lobule_R | 70 | SM network |
| Heschl_L | 79 | SM network |
| Heschl_R | 80 | SM network |
| Temporal_Sup_L | 81 | SM network |
| Temporal_Sup_R | 82 | SM network |
| Parietal_Sup_L | 59 | DAN |
| Parietal_Sup_R | 60 | DAN |
| Insula_L | 29 | VAN |

| | | |
|---|---|---|
| Insula_R | 30 | VAN |
| SupraMarginal_L | 63 | VAN |
| SupraMarginal_R | 64 | VAN |
| Frontal_Sup_Orb_L | 5 | Limbic network |
| Frontal_Sup_Orb_R | 6 | Limbic network |
| Olfactory_L | 21 | Limbic network |
| Olfactory_R | 22 | Limbic network |
| Rectus_L | 27 | Limbic network |
| Rectus_R | 28 | Limbic network |
| Hippocampus_L | 37 | Limbic network |
| Hippocampus_R | 38 | Limbic network |
| ParaHippocampal_L | 39 | Limbic network |
| ParaHippocampal_R | 40 | Limbic network |
| Amygdala_L | 41 | Limbic network |
| Amygdala_R | 42 | Limbic network |
| Temporal_Pole_Sup_L | 83 | Limbic network |
| Temporal_Pole_Sup_R | 84 | Limbic network |
| Temporal_Pole_Mid_L | 87 | Limbic network |
| Temporal_Pole_Mid_R | 88 | Limbic network |
| Temporal_Inf_L | 89 | Limbic network |
| Temporal_Inf_R | 90 | Limbic network |
| Frontal_Mid_L | 7 | FTP Cognitive network |
| Frontal_Mid_R | 8 | FTP Cognitive network |
| Frontal_Mid_Orb_L | 9 | FTP Cognitive network |
| Frontal_Mid_Orb_R | 10 | FTP Cognitive network |
| Frontal_Inf_Oper_L | 11 | FTP Cognitive network |
| Frontal_Inf_Oper_R | 12 | FTP Cognitive network |
| Frontal_Inf_Tri_L | 13 | FTP Cognitive network |
| Frontal_Inf_Tri_R | 14 | FTP Cognitive network |
| Cingulum_Post_L | 35 | FTP Cognitive network |
| Cingulum_Post_R | 36 | FTP Cognitive network |
| Parietal_Inf_L | 61 | FTP Cognitive network |
| Parietal_Inf_R | 62 | FTP Cognitive network |
| Frontal_Sup_L | 3 | DMN |
| Frontal_Sup_R | 4 | DMN |
| Frontal_Inf_Orb_L | 15 | DMN |
| Frontal_Inf_Orb_R | 16 | DMN |
| Frontal_Sup_Medial_L | 23 | DMN |
| Frontal_Sup_Medial_R | 24 | DMN |
| Frontal_Med_Orb_L | 25 | DMN |
| Frontal_Med_Orb_R | 26 | DMN |
| Cingulum_Ant_L | 31 | DMN |
| Cingulum_Ant_R | 32 | DMN |
| Angular_L | 65 | DMN |
| Angular_R | 66 | DMN |
| Precuneus_L | 67 | DMN |
| Precuneus_R | 68 | DMN |
| Temporal_Mid_L | 85 | DMN |
| Temporal_Mid_R | 86 | DMN |

| | | |
|---|---|---|
| Caudate_L | 71 | Basal gansalia network |
| Caudate_R | 72 | Basal gansalia network |
| Putamen_L | 73 | Basal gansalia network |
| Putamen_R | 74 | Basal gansalia network |
| Pallidum_L | 75 | Basal gansalia network |
| Pallidum_R | 76 | Basal gansalia network |
| Thalamus_L | 77 | Basal gansalia network |
| Thalamus_R | 78 | Basal gansalia network |
| Cerebelum_Crus1_L | 91 | Cerebellum |
| Cerebelum_Crus1_R | 92 | Cerebellum |
| Cerebelum_Crus2_L | 93 | Cerebellum |
| Cerebelum_Crus2_R | 94 | Cerebellum |
| Cerebelum_3_L | 95 | Cerebellum |
| Cerebelum_3_R | 96 | Cerebellum |
| Cerebelum_4_5_L | 97 | Cerebellum |
| Cerebelum_4_5_R | 98 | Cerebellum |
| Cerebelum_6_L | 99 | Cerebellum |
| Cerebelum_6_R | 100 | Cerebellum |
| Cerebelum_7b_L | 101 | Cerebellum |
| Cerebelum_7b_R | 102 | Cerebellum |
| Cerebelum_8_L | 103 | Cerebellum |
| Cerebelum_8_R | 104 | Cerebellum |
| Cerebelum_9_L | 105 | Cerebellum |
| Cerebelum_9_R | 106 | Cerebellum |
| Cerebelum_10_L | 107 | Cerebellum |
| Cerebelum_10_R | 108 | Cerebellum |
| Vermis_1_2 | 109 | Cerebellum |
| Vermis_3 | 110 | Cerebellum |
| Vermis_4_5 | 111 | Cerebellum |
| Vermis_6 | 112 | Cerebellum |
| Vermis_7 | 113 | Cerebellum |
| Vermis_8 | 114 | Cerebellum |
| Vermis_9 | 115 | Cerebellum |
| Vermis_10 | 116 | Cerebellum |

SM: sensori-motor area; DAN: dorsal attention network; VAN: ventral attention network; FTP: frontal-temporal-parietal; DMN: default mode network.

**Table S2.** Functional network mapping of jointed atlas into nine networks

| | Region Name | ROI Label | ROI Name |
|---|---|---|---|
| **Schaefer's atlas** | | | |
| | Vis_1 | 1 | Visual network |
| | Vis_2 | 2 | Visual network |
| | Vis_3 | 3 | Visual network |
| | Vis_4 | 4 | Visual network |
| | Vis_5 | 5 | Visual network |
| | Vis_6 | 6 | Visual network |
| | Vis_7 | 7 | Visual network |
| | Vis_8 | 8 | Visual network |
| | Vis_9 | 9 | Visual network |
| | Vis_10 | 10 | Visual network |
| | Vis_11 | 11 | Visual network |
| | Vis_12 | 12 | Visual network |
| | Vis_13 | 13 | Visual network |
| | Vis_14 | 14 | Visual network |
| | Vis_15 | 15 | Visual network |
| | Vis_16 | 16 | Visual network |
| | Vis_17 | 17 | Visual network |
| | Vis_18 | 18 | Visual network |
| | Vis_19 | 19 | Visual network |
| | Vis_20 | 20 | Visual network |
| | Vis_21 | 21 | Visual network |
| | Vis_22 | 22 | Visual network |
| | Vis_23 | 23 | Visual network |
| | Vis_24 | 24 | Visual network |
| | Vis_25 | 25 | Visual network |
| | Vis_26 | 26 | Visual network |
| | Vis_27 | 27 | Visual network |
| | Vis_28 | 28 | Visual network |
| | Vis_29 | 29 | Visual network |
| | Vis_30 | 30 | Visual network |
| | Vis_31 | 31 | Visual network |
| | SomMot_1 | 32 | SM network |
| | SomMot_2 | 33 | SM network |
| | SomMot_3 | 34 | SM network |
| | SomMot_4 | 35 | SM network |
| | SomMot_5 | 36 | SM network |
| | SomMot_6 | 37 | SM network |
| | SomMot_7 | 38 | SM network |
| | SomMot_8 | 39 | SM network |
| | SomMot_9 | 40 | SM network |

| | | |
|---|---|---|
| SomMot_10 | 41 | SM network |
| SomMot_11 | 42 | SM network |
| SomMot_12 | 43 | SM network |
| SomMot_13 | 44 | SM network |
| SomMot_14 | 45 | SM network |
| SomMot_15 | 46 | SM network |
| SomMot_16 | 47 | SM network |
| SomMot_17 | 48 | SM network |
| SomMot_18 | 49 | SM network |
| SomMot_19 | 50 | SM network |
| SomMot_20 | 51 | SM network |
| SomMot_21 | 52 | SM network |
| SomMot_22 | 53 | SM network |
| SomMot_23 | 54 | SM network |
| SomMot_24 | 55 | SM network |
| SomMot_25 | 56 | SM network |
| SomMot_26 | 57 | SM network |
| SomMot_27 | 58 | SM network |
| SomMot_28 | 59 | SM network |
| SomMot_29 | 60 | SM network |
| SomMot_30 | 61 | SM network |
| SomMot_31 | 62 | SM network |
| SomMot_32 | 63 | SM network |
| SomMot_33 | 64 | SM network |
| SomMot_34 | 65 | SM network |
| SomMot_35 | 66 | SM network |
| SomMot_36 | 67 | SM network |
| SomMot_37 | 68 | SM network |
| Post_1 | 69 | DAN |
| Post_2 | 70 | DAN |
| Post_3 | 71 | DAN |
| Post_4 | 72 | DAN |
| Post_5 | 73 | DAN |
| Post_6 | 74 | DAN |
| Post_7 | 75 | DAN |
| Post_8 | 76 | DAN |
| Post_9 | 77 | DAN |
| Post_10 | 78 | DAN |
| Post_11 | 79 | DAN |
| Post_12 | 80 | DAN |
| Post_13 | 81 | DAN |
| Post_14 | 82 | DAN |
| Post_15 | 83 | DAN |
| Post_16 | 84 | DAN |
| Post_17 | 85 | DAN |

| | | |
|---|---|---|
| FEF_1 | 86 | DAN |
| FEF_2 | 87 | DAN |
| FEF_3 | 88 | DAN |
| FEF_4 | 89 | DAN |
| PrCv_1 | 90 | DAN |
| PrCv_2 | 91 | DAN |
| ParOper_1 | 92 | VAN |
| ParOper_2 | 93 | VAN |
| ParOper_3 | 94 | VAN |
| ParOper_4 | 95 | VAN |
| TempOcc_1 | 96 | VAN |
| FrOperIns_1 | 97 | VAN |
| FrOperIns_2 | 98 | VAN |
| FrOperIns_3 | 99 | VAN |
| FrOperIns_4 | 100 | VAN |
| FrOperIns_5 | 101 | VAN |
| FrOperIns_6 | 102 | VAN |
| FrOperIns_7 | 103 | VAN |
| FrOperIns_8 | 104 | VAN |
| FrOperIns_9 | 105 | VAN |
| PFCl_1 | 106 | VAN |
| Med_1 | 107 | VAN |
| Med_2 | 108 | VAN |
| Med_3 | 109 | VAN |
| Med_4 | 110 | VAN |
| Med_5 | 111 | VAN |
| Med_6 | 112 | VAN |
| Med_7 | 113 | VAN |
| OFC_1 | 114 | Limbic network |
| OFC_2 | 115 | Limbic network |
| OFC_3 | 116 | Limbic network |
| OFC_4 | 117 | Limbic network |
| OFC_5 | 118 | Limbic network |
| TempPole_1 | 119 | Limbic network |
| TempPole_2 | 120 | Limbic network |
| TempPole_3 | 121 | Limbic network |
| TempPole_4 | 122 | Limbic network |
| TempPole_5 | 123 | Limbic network |
| TempPole_6 | 124 | Limbic network |
| TempPole_7 | 125 | Limbic network |
| TempPole_8 | 126 | Limbic network |
| Par_1 | 127 | FTP Cognitive network |
| Par_2 | 128 | FTP Cognitive network |
| Par_3 | 129 | FTP Cognitive network |
| Par_4 | 130 | FTP Cognitive network |

| | | |
|---|---|---|
| Par_5 | 131 | FTP Cognitive network |
| Par_6 | 132 | FTP Cognitive network |
| Temp_1 | 133 | FTP Cognitive network |
| OFC_1 | 134 | FTP Cognitive network |
| PFCl_1 | 135 | FTP Cognitive network |
| PFCl_2 | 136 | FTP Cognitive network |
| PFCl_3 | 137 | FTP Cognitive network |
| PFCl_4 | 138 | FTP Cognitive network |
| PFCl_5 | 139 | FTP Cognitive network |
| PFCl_6 | 140 | FTP Cognitive network |
| PFCl_7 | 141 | FTP Cognitive network |
| PFCl_8 | 142 | FTP Cognitive network |
| PFCv_1 | 143 | FTP Cognitive network |
| pCun_1 | 144 | FTP Cognitive network |
| pCun_2 | 145 | FTP Cognitive network |
| Cing_1 | 146 | FTP Cognitive network |
| Cing_2 | 147 | FTP Cognitive network |
| PFCmp_1 | 148 | FTP Cognitive network |
| Temp_1 | 149 | DMN |
| Temp_2 | 150 | DMN |
| Temp_3 | 151 | DMN |
| Temp_4 | 152 | DMN |
| Temp_5 | 153 | DMN |
| Temp_6 | 154 | DMN |
| Temp_7 | 155 | DMN |
| Temp_8 | 156 | DMN |
| Temp_9 | 157 | DMN |
| Temp_10 | 158 | DMN |
| Par_1 | 159 | DMN |
| Par_2 | 160 | DMN |
| Par_3 | 161 | DMN |
| Par_4 | 162 | DMN |
| Par_5 | 163 | DMN |
| Par_6 | 164 | DMN |
| Par_7 | 165 | DMN |
| PFC_1 | 166 | DMN |
| PFC_2 | 167 | DMN |
| PFC_3 | 168 | DMN |
| PFC_4 | 169 | DMN |
| PFC_5 | 170 | DMN |
| PFC_6 | 171 | DMN |
| PFC_7 | 172 | DMN |
| PFC_8 | 173 | DMN |
| PFC_9 | 174 | DMN |
| PFC_10 | 175 | DMN |

| | | |
|---|---|---|
| PFC_11 | 176 | DMN |
| PFC_12 | 177 | DMN |
| PFC_13 | 178 | DMN |
| PFC_14 | 179 | DMN |
| PFC_15 | 180 | DMN |
| PFC_16 | 181 | DMN |
| PFC_17 | 182 | DMN |
| PFC_18 | 183 | DMN |
| PFC_19 | 184 | DMN |
| PFC_20 | 185 | DMN |
| PFC_21 | 186 | DMN |
| PFC_22 | 187 | DMN |
| PFC_23 | 188 | DMN |
| PFC_24 | 189 | DMN |
| pCunPCC_1 | 190 | DMN |
| pCunPCC_2 | 191 | DMN |
| pCunPCC_3 | 192 | DMN |
| pCunPCC_4 | 193 | DMN |
| pCunPCC_5 | 194 | DMN |
| pCunPCC_6 | 195 | DMN |
| pCunPCC_7 | 196 | DMN |
| pCunPCC_8 | 197 | DMN |
| pCunPCC_9 | 198 | DMN |
| pCunPCC_10 | 199 | DMN |
| pCunPCC_11 | 200 | DMN |
| Vis_1 | 201 | Visual network |
| Vis_2 | 202 | Visual network |
| Vis_3 | 203 | Visual network |
| Vis_4 | 204 | Visual network |
| Vis_5 | 205 | Visual network |
| Vis_6 | 206 | Visual network |
| Vis_7 | 207 | Visual network |
| Vis_8 | 208 | Visual network |
| Vis_9 | 209 | Visual network |
| Vis_10 | 210 | Visual network |
| Vis_11 | 211 | Visual network |
| Vis_12 | 212 | Visual network |
| Vis_13 | 213 | Visual network |
| Vis_14 | 214 | Visual network |
| Vis_15 | 215 | Visual network |
| Vis_16 | 216 | Visual network |
| Vis_17 | 217 | Visual network |
| Vis_18 | 218 | Visual network |
| Vis_19 | 219 | Visual network |
| Vis_20 | 220 | Visual network |

| | | |
|---|---|---|
| Vis_21 | 221 | Visual network |
| Vis_22 | 222 | Visual network |
| Vis_23 | 223 | Visual network |
| Vis_24 | 224 | Visual network |
| Vis_25 | 225 | Visual network |
| Vis_26 | 226 | Visual network |
| Vis_27 | 227 | Visual network |
| Vis_28 | 228 | Visual network |
| Vis_29 | 229 | Visual network |
| Vis_30 | 230 | Visual network |
| SomMot_1 | 231 | SM network |
| SomMot_2 | 232 | SM network |
| SomMot_3 | 233 | SM network |
| SomMot_4 | 234 | SM network |
| SomMot_5 | 235 | SM network |
| SomMot_6 | 236 | SM network |
| SomMot_7 | 237 | SM network |
| SomMot_8 | 238 | SM network |
| SomMot_9 | 239 | SM network |
| SomMot_10 | 240 | SM network |
| SomMot_11 | 241 | SM network |
| SomMot_12 | 242 | SM network |
| SomMot_13 | 243 | SM network |
| SomMot_14 | 244 | SM network |
| SomMot_15 | 245 | SM network |
| SomMot_16 | 246 | SM network |
| SomMot_17 | 247 | SM network |
| SomMot_18 | 248 | SM network |
| SomMot_19 | 249 | SM network |
| SomMot_20 | 250 | SM network |
| SomMot_21 | 251 | SM network |
| SomMot_22 | 252 | SM network |
| SomMot_23 | 253 | SM network |
| SomMot_24 | 254 | SM network |
| SomMot_25 | 255 | SM network |
| SomMot_26 | 256 | SM network |
| SomMot_27 | 257 | SM network |
| SomMot_28 | 258 | SM network |
| SomMot_29 | 259 | SM network |
| SomMot_30 | 260 | SM network |
| SomMot_31 | 261 | SM network |
| SomMot_32 | 262 | SM network |
| SomMot_33 | 263 | SM network |
| SomMot_34 | 264 | SM network |
| SomMot_35 | 265 | SM network |

| | | |
|---|---|---|
| SomMot_36 | 266 | SM network |
| SomMot_37 | 267 | SM network |
| SomMot_38 | 268 | SM network |
| SomMot_39 | 269 | SM network |
| SomMot_40 | 270 | SM network |
| Post_1 | 271 | DAN |
| Post_2 | 272 | DAN |
| Post_3 | 273 | DAN |
| Post_4 | 274 | DAN |
| Post_5 | 275 | DAN |
| Post_6 | 276 | DAN |
| Post_7 | 277 | DAN |
| Post_8 | 278 | DAN |
| Post_9 | 279 | DAN |
| Post_10 | 280 | DAN |
| Post_11 | 281 | DAN |
| Post_12 | 282 | DAN |
| Post_13 | 283 | DAN |
| Post_14 | 284 | DAN |
| Post_15 | 285 | DAN |
| Post_16 | 286 | DAN |
| Post_17 | 287 | DAN |
| Post_18 | 288 | DAN |
| Post_19 | 289 | DAN |
| FEF_1 | 290 | DAN |
| FEF_2 | 291 | DAN |
| FEF_3 | 292 | DAN |
| PrCv_1 | 293 | DAN |
| TempOccPar_1 | 294 | VAN |
| TempOccPar_2 | 295 | VAN |
| TempOccPar_3 | 296 | VAN |
| TempOccPar_4 | 297 | VAN |
| TempOccPar_5 | 298 | VAN |
| TempOccPar_6 | 299 | VAN |
| TempOccPar_7 | 300 | VAN |
| PrC_1 | 301 | VAN |
| FrOperIns_1 | 302 | VAN |
| FrOperIns_2 | 303 | VAN |
| FrOperIns_3 | 304 | VAN |
| FrOperIns_4 | 305 | VAN |
| FrOperIns_5 | 306 | VAN |
| FrOperIns_6 | 307 | VAN |
| FrOperIns_7 | 308 | VAN |
| FrOperIns_8 | 309 | VAN |
| PFCl_1 | 310 | VAN |

| | | |
|---|---|---|
| Med_1 | 311 | VAN |
| Med_2 | 312 | VAN |
| Med_3 | 313 | VAN |
| Med_4 | 314 | VAN |
| Med_5 | 315 | VAN |
| Med_6 | 316 | VAN |
| Med_7 | 317 | VAN |
| Med_8 | 318 | VAN |
| OFC_1 | 319 | Limbic network |
| OFC_2 | 320 | Limbic network |
| OFC_3 | 321 | Limbic network |
| OFC_4 | 322 | Limbic network |
| OFC_5 | 323 | Limbic network |
| OFC_6 | 324 | Limbic network |
| TempPole_1 | 325 | Limbic network |
| TempPole_2 | 326 | Limbic network |
| TempPole_3 | 327 | Limbic network |
| TempPole_4 | 328 | Limbic network |
| TempPole_5 | 329 | Limbic network |
| TempPole_6 | 330 | Limbic network |
| TempPole_7 | 331 | Limbic network |
| Par_1 | 332 | FTP Cognitive network |
| Par_2 | 333 | FTP Cognitive network |
| Par_3 | 334 | FTP Cognitive network |
| Par_4 | 335 | FTP Cognitive network |
| Par_5 | 336 | FTP Cognitive network |
| Par_6 | 337 | FTP Cognitive network |
| Temp_1 | 338 | FTP Cognitive network |
| Temp_2 | 339 | FTP Cognitive network |
| PFCv_1 | 340 | FTP Cognitive network |
| PFCl_1 | 341 | FTP Cognitive network |
| PFCl_2 | 342 | FTP Cognitive network |
| PFCl_3 | 343 | FTP Cognitive network |
| PFCl_4 | 344 | FTP Cognitive network |
| PFCl_5 | 345 | FTP Cognitive network |
| PFCl_6 | 346 | FTP Cognitive network |
| PFCl_7 | 347 | FTP Cognitive network |
| PFCl_8 | 348 | FTP Cognitive network |
| PFCl_9 | 349 | FTP Cognitive network |
| PFCl_10 | 350 | FTP Cognitive network |
| PFCl_11 | 351 | FTP Cognitive network |
| PFCl_12 | 352 | FTP Cognitive network |
| PFCl_13 | 353 | FTP Cognitive network |
| PFCl_14 | 354 | FTP Cognitive network |
| PFCl_15 | 355 | FTP Cognitive network |

| | | |
|---|---|---|
| pCun_1 | 356 | FTP Cognitive network |
| pCun_2 | 357 | FTP Cognitive network |
| Cing_1 | 358 | FTP Cognitive network |
| Cing_2 | 359 | FTP Cognitive network |
| PFCmp_1 | 360 | FTP Cognitive network |
| PFCmp_2 | 361 | FTP Cognitive network |
| Par_1 | 362 | DMN |
| Par_2 | 363 | DMN |
| Par_3 | 364 | DMN |
| Par_4 | 365 | DMN |
| Par_5 | 366 | DMN |
| Temp_1 | 367 | DMN |
| Temp_2 | 368 | DMN |
| Temp_3 | 369 | DMN |
| Temp_4 | 370 | DMN |
| Temp_5 | 371 | DMN |
| Temp_6 | 372 | DMN |
| Temp_7 | 373 | DMN |
| Temp_8 | 374 | DMN |
| PFCv_1 | 375 | DMN |
| PFCv_2 | 376 | DMN |
| PFCv_3 | 377 | DMN |
| PFCv_4 | 378 | DMN |
| PFCdPFCm_1 | 379 | DMN |
| PFCdPFCm_2 | 380 | DMN |
| PFCdPFCm_3 | 381 | DMN |
| PFCdPFCm_4 | 382 | DMN |
| PFCdPFCm_5 | 383 | DMN |
| PFCdPFCm_6 | 384 | DMN |
| PFCdPFCm_7 | 385 | DMN |
| PFCdPFCm_8 | 386 | DMN |
| PFCdPFCm_9 | 387 | DMN |
| PFCdPFCm_10 | 388 | DMN |
| PFCdPFCm_11 | 389 | DMN |
| PFCdPFCm_12 | 390 | DMN |
| PFCdPFCm_13 | 391 | DMN |
| pCunPCC_1 | 392 | DMN |
| pCunPCC_2 | 393 | DMN |
| pCunPCC_3 | 394 | DMN |
| pCunPCC_4 | 395 | DMN |
| pCunPCC_5 | 396 | DMN |
| pCunPCC_6 | 397 | DMN |
| pCunPCC_7 | 398 | DMN |
| pCunPCC_8 | 399 | DMN |
| pCunPCC_9 | 400 | DMN |

| | | |
|---|---|---|
| Thalamus_L | 4 | Basal gansalia network |
| Caudate_L | 5 | Basal gansalia network |
| Putamen_L | 6 | Basal gansalia network |
| Pallidum_L | 7 | Basal gansalia network |
| Hippocampus_L | 9 | Basal gansalia network |
| Amygdala_L | 10 | Basal gansalia network |
| Accumbens_L | 11 | Basal gansalia network |
| Thalamus_R | 15 | Basal gansalia network |
| Caudate_R | 16 | Basal gansalia network |
| Putamen_R | 17 | Basal gansalia network |
| Pallidum_R | 18 | Basal gansalia network |
| Hippocampus_R | 19 | Basal gansalia network |
| Amygdala_R | 20 | Basal gansalia network |
| Accumbens_R | 21 | Basal gansalia network |
| Left_I_IV | 1 | Cerebellum |
| Right_I_IV | 2 | Cerebellum |
| Left_V | 3 | Cerebellum |
| Right_V | 4 | Cerebellum |
| Left_VI | 5 | Cerebellum |
| Vermis_VI | 6 | Cerebellum |
| Right_VI | 7 | Cerebellum |
| Left_CrusI | 8 | Cerebellum |
| Vermis_CrusI | 9 | Cerebellum |
| Right_CrusI | 10 | Cerebellum |
| Left_CrusII | 11 | Cerebellum |
| Vermis_CrusII | 12 | Cerebellum |
| Right_CrusII | 13 | Cerebellum |
| Left_VIIb | 14 | Cerebellum |
| Vermis_VIIb | 15 | Cerebellum |
| Right_VIIb | 16 | Cerebellum |
| Left_VIIIa | 17 | Cerebellum |
| Vermis_VIIIa | 18 | Cerebellum |
| Right_VIIIa | 19 | Cerebellum |
| Left_VIIIb | 20 | Cerebellum |
| Vermis_VIIIb | 21 | Cerebellum |
| Right_VIIIb | 22 | Cerebellum |
| Left_IX | 23 | Cerebellum |
| Vermis_IX | 24 | Cerebellum |
| Right_IX | 25 | Cerebellum |
| Left_X | 26 | Cerebellum |
| Vermis_X | 27 | Cerebellum |
| Right_X | 28 | Cerebellum |

SM: sensori-motor area; DAN: dorsal attention network; VAN: ventral attention network; FTP: frontal-temporal-parietal; DMN: default mode network.

**Table S3.** Overlap information between different atlases

| ROI label in HO | Voxel number in HO | ROI label in Schaefer | Voxel number in Schaefer | Voxel number of overlap |
|---|---|---|---|---|
| **Subcortical HO & cortex Schaefer** | | | | |
| 4 | 1470 | 13 | 119 | 1 |
| 9 | 765 | 2 | 425 | 8 |
| 9 | 765 | 7 | 183 | 2 |
| 9 | 765 | 20 | 120 | 2 |
| 9 | 765 | 126 | 203 | 7 |
| 10 | 332 | 125 | 431 | 2 |
| 11 | 89 | 117 | 405 | 2 |
| 15 | 1401 | 214 | 217 | 1 |
| 19 | 765 | 202 | 348 | 2 |
| 19 | 765 | 214 | 217 | 1 |
| 19 | 765 | 217 | 127 | 1 |
| 19 | 765 | 328 | 481 | 1 |
| 19 | 765 | 331 | 279 | 9 |
| 20 | 413 | 329 | 471 | 12 |
| 21 | 84 | 323 | 309 | 1 |
| **Subcortical HO & cerebellum SUIT** | | | | |
| ROI label in HO | Voxel number in HO | ROI label in SUIT | Voxel number in SUIT | Voxel number of overlap |
| 4 | 1470 | 1 | 935 | 1 |
| **Cerebellum SUIT & cortex Schaefer** | | | | |
| ROI label in HO | Voxel number in HO | ROI label in Schaefer | Voxel number in Schaefer | Voxel number of overlap |
| 1 | 935 | 2 | 425 | 3 |
| 1 | 935 | 7 | 183 | 19 |
| 1 | 935 | 10 | 261 | 3 |
| 1 | 935 | 13 | 119 | 18 |
| 1 | 935 | 126 | 203 | 5 |
| 2 | 1004 | 201 | 363 | 9 |
| 2 | 1004 | 202 | 348 | 11 |
| 2 | 1004 | 207 | 136 | 22 |
| 2 | 1004 | 212 | 348 | 15 |
| 2 | 1004 | 214 | 217 | 4 |
| 2 | 1004 | 331 | 279 | 3 |
| 3 | 978 | 1 | 347 | 51 |
| 3 | 978 | 2 | 425 | 24 |
| 3 | 978 | 4 | 401 | 12 |
| 3 | 978 | 7 | 183 | 21 |
| 3 | 978 | 10 | 261 | 54 |
| 3 | 978 | 13 | 119 | 10 |
| 3 | 978 | 126 | 203 | 2 |
| 4 | 932 | 201 | 363 | 69 |

| | | | | |
|---|---|---|---|---|
| 4 | 932 | 202 | 348 | 5 |
| 4 | 932 | 203 | 355 | 8 |
| 4 | 932 | 205 | 453 | 26 |
| 4 | 932 | 207 | 136 | 11 |
| 4 | 932 | 211 | 328 | 2 |
| 4 | 932 | 212 | 348 | 57 |
| 5 | 1915 | 1 | 347 | 85 |
| 5 | 1915 | 3 | 309 | 40 |
| 5 | 1915 | 4 | 401 | 19 |
| 5 | 1915 | 5 | 426 | 8 |
| 5 | 1915 | 9 | 320 | 5 |
| 5 | 1915 | 10 | 261 | 10 |
| 5 | 1915 | 11 | 308 | 8 |
| 5 | 1915 | 69 | 363 | 9 |
| 7 | 1772 | 201 | 363 | 33 |
| 7 | 1772 | 203 | 355 | 75 |
| 7 | 1772 | 204 | 386 | 9 |
| 7 | 1772 | 205 | 453 | 13 |
| 7 | 1772 | 206 | 530 | 35 |
| 7 | 1772 | 208 | 279 | 1 |
| 7 | 1772 | 211 | 328 | 24 |
| 7 | 1772 | 271 | 738 | 1 |
| 8 | 2716 | 3 | 309 | 4 |
| 8 | 2716 | 69 | 363 | 25 |
| 8 | 2716 | 70 | 612 | 9 |
| 10 | 2767 | 203 | 355 | 2 |
| 10 | 2767 | 204 | 386 | 9 |
| 10 | 2767 | 208 | 279 | 2 |
| 10 | 2767 | 215 | 719 | 1 |
| 10 | 2767 | 271 | 738 | 13 |
| 13 | 1998 | 330 | 434 | 1 |

**Table S4.** Graph theoretical measures of clustering results based on 16-min data. Btsp.: bootstrapping; C.I.: confidence interval; Ses.: session.

| | Not-centered | | | centered (adding mean back) | | |
|---|---|---|---|---|---|---|
| **AAL atlas (16 min)** | | | | | | |
| **State S** | Btsp. C.I. of ses. A | ses. A | ses. B | Btsp. C.I. of ses. A | ses. A | ses. B |
| global efficiency | [0.3207, 0.3500] | 0.3297 | 0.3316 | [0.3243, 0.3726] | 0.3425 | 0.3308 |
| modularity | [0.1465, 0.1880] | 0.1607 | 0.1814 | [0.1158, 0.1598] | 0.1358 | 0.1528 |
| **State I** | | | | | | |
| global efficiency | [0.4855, 0.5516] | 0.5096 | 0.5031 | [0.5063, 0.5972] | 0.5473 | 0.5272 |
| modularity | [0.0577, 0.0794] | 0.0702 | 0.0763 | [0.0540, 0.0740] | 0.0637 | 0.0736 |
| **Joint atlas (16 min)** | | | | | | |
| **State S** | Btsp. C.I. of ses. A | ses. A | ses. B | Btsp. C.I. of ses. A | ses. A | ses. B |
| global efficiency | [0.2860, 0.3182] | 0.2933 | 0.3028 | [0.2956, 0.3464] | 0.3125 | 0.3070 |
| modularity | [0.1796, 0.2072] | 0.1904 | 0.2009 | [0.1218, 0.1659] | 0.1433 | 0.1553 |
| **State I** | | | | | | |
| global efficiency | [0.4440, 0.5027] | 0.4660 | 0.4622 | [0.4736, 0.5696] | 0.5170 | 0.5022 |
| modularity | [0.0589, 0.0722] | 0.0639 | 0.0707 | [0.0511, 0.0694] | 0.0581 | 0.0681 |

**Table S5.** Available subjects for the parameters of mean dwell time (MDT) and inter-transition interval (ITI)

| **Centralization** | **Time length** | **ROIs** | **Session** | **MDT of state S** | **MDT of state I** | **ITI** |
|---|---|---|---|---|---|---|
| No | 16 min. | 116 | A | 20 | 20 | 20 |
| | 16 min. | 116 | B | 22 | 21 | 21 |
| No | 8 min. | 116 | A | 19 | 20 | 19 |
| | 8 min. | 116 | B | 21 | 22 | 21 |
| No | 16 min. | 442 | A | 20 | 20 | 20 |
| | 16 min. | 442 | B | 21 | 22 | 21 |
| No | 8 min. | 442 | A | 16 | 17 | 15 |
| | 8 min. | 442 | B | 21 | 20 | 20 |
| Yes | 16 min. | 116 | A | 23 | 23 | 23 |
| | 16 min. | 116 | B | 23 | 23 | 23 |
| Yes | 8 min. | 116 | A | 23 | 23 | 23 |
| | 8 min. | 116 | B | 23 | 23 | 23 |
| Yes | 16 min. | 442 | A | 23 | 23 | 23 |
| | 16 min. | 442 | B | 23 | 23 | 23 |
| Yes | 8 min. | 442 | A | 23 | 22 | 22 |
| | 8 min. | 442 | B | 23 | 23 | 23 |

**Table S6.** Cosine distance between the 2 clusters for the two sessions

**a) Clusters based on not-centered data**

| | | AAL atlas | | | | Joint atlas | | | |
|---|---|---|---|---|---|---|---|---|---|
| | | **16-min data** | | **8-min data** | | **16-min data** | | **8-min data** | |
| | | **Session B** | | **Session B** | | **Session B** | | **Session B** | |
| | | State S | State I | State S | State I | State S | State I | State S | State I |
| **Session A** | State S | 0.0132 | 0.0917 | 0.0218 | 0.0967 | 0.0199 | 0.1260 | 0.0332 | 0.1284 |
| | State I | 0.1070 | 0.0039 | 0.1158 | 0.0068 | 0.1404 | 0.0044 | 0.1479 | 0.0083 |

**b) Clusters based on centering data**

| | | AAL atlas | | | | Joint atlas | | | |
|---|---|---|---|---|---|---|---|---|---|
| | | **16-min data** | | **8-min data** | | **16-min data** | | **8-min data** | |
| | | **Session B** | | **Session B** | | **Session B** | | **Session B** | |
| | | State S | State I | State S | State I | State S | State I | State S | State I |
| **Session A** | State S | 0.0137 | 1.9861 | 0.0292 | 1.9699 | 0.0162 | 1.9828 | 0.0380 | 1.9602 |
| | State I | 1.9858 | 0.0141 | 1.9705 | 0.0303 | 1.9836 | 0.0172 | 1.9612 | 0.0406 |

**c) Clusters based on not-centered data & removing session mean matrices**

| | | AAL atlas | | | | Joint atlas | | | |
|---|---|---|---|---|---|---|---|---|---|
| | | **16 min data** | | **8 min data** | | **16 min data** | | **8 min data** | |
| | | **Session B** | | **Session B** | | **Session B** | | **Session B** | |
| | | State S | State I | State S | State I | State S | State I | State S | State I |
| **Session A** | State S | 0.0444 | 1.9556 | 0.0769 | 1.9231 | 0.0427 | 1.9573 | 0.0839 | 1.9161 |
| | State I | 1.9556 | 0.0444 | 1.9231 | 0.0769 | 1.9573 | 0.0427 | 1.9161 | 0.0839 |

**Table S7.** Reproducibility of DFC parameters in terms of Spearman's correlations . CI: confidence interval based on bootstrapping (1000 times). Bold: significant reliability.

| | | 116 *R*OIs | | | | 442 *R*OIs | | | |
|---|---|---|---|---|---|---|---|---|---|
| | | 16-min data | | 8-min data | | 16-min data | | 8-min data | |
| | | **Not-centered** | **Centered** | **Not-centered** | **Centered** | **Not-centered** | **Centered** | **Not-centered** | **Centered** |
| **Mean dwell time** | **state S** | *rho* = 0.05 | *rho* = 0.05 | *rho* = -0.02 | *rho* = 0.30 | *rho* = -0.40 | *rho* = -0.13 | *rho* = -0.04 | *rho* = 0.39 |
| | | *p* = 0.8372 | *p* = 0.8124 | *p* = 0.9410 | *p* = 0.1593 | *p* = 0.0849 | *p* = 0.5485 | *p* = 0.8828 | *p* = 0.0703 |
| | | CI: [-0.4303, 0.4956] | CI: [-0.4762, 0.5155] | CI: [-0.4886, 0.4305] | CI: [-0.1756, 0.6996] | CI: [-0.7058, -0.0004] | CI: [-0.5734, 0.3784] | CI: [-0.6159, 0.5077] | CI: [-0.0972, 0.7602] |
| | **state I** | *rho* = 0.13 | *rho* = 0.22 | *rho* = 0.21 | *rho* = -0.00 | *rho* = -0.08 | *rho* = 0.34 | *rho* = 0.18 | *rho* = 0.34 |
| | | *p* = 0.5852 | *p* = 0.3064 | *p* = 0.4109 | *p* = 0.9874 | *p* = 0.7482 | *p* = 0.1158 | *p* = 0.5235 | *p* = 0.1124 |
| | | CI: [-0.3707, 0.6225] | CI: [-0.2728, 0.6107] | CI: [-0.3559, 0.6882] | CI: [-0.4301, 0.4245] | CI: [-0.6119, 0.4470] | CI: [0.0025, 0.5945] | CI: [-0.3559, 0.6885] | CI: [-0.1370, 0.7236] |
| **Prevalence** | **state S** | ***rho* = 0.53** | *rho* = 0.08 | *rho* = 0.32 | *rho* = 0.20 | ***rho* = 0.44** | *rho* = 0.32 | *rho* = 0.31 | *rho* = -0.04 |
| | | ***p* = 0.0097** | *p* = 0.7329 | *p* = 0.1356 | *p* = 0.3698 | ***p* = 0.0341** | *p* = 0.1380 | *p* = 0.1460 | *p* = 0.8718 |
| | | **CI: [0.1337, 0.7943]** | CI: [-0.3712, 0.5249] | CI: [-0.1220, 0.6384] | CI: [-0.2155, 0.5733] | **CI: [-0.0443, 0.7576]** | CI: [-0.1322, 0.6704] | CI: [-0.1460, 0.6771] | CI: [-0.4778, 0.4745] |
| **Inter-transition interval** | | *rho* = -0.12 | *rho* = 0.09 | *rho* = 0.20 | *rho* = 0.29 | *rho* = -0.26 | *rho* = -0.33 | *rho* = 0.35 | *rho* = 0.25 |
| | | *p* = 0.6104 | *p* = 0.6659 | *p* = 0.4377 | *p* = 0.1722 | *p* = 0.2719 | *p* = 0.1257 | *p* = 0.2271 | *p* = 0.2582 |
| | | CI: [-0.5725, 0.3182] | CI: [-0.4053, 0.5805] | CI: [-0.3743, 0.7433] | CI: [-0.1707, 0.6872] | CI: [-0.6270, 0.1902] | CI: [-0.7364, 0.2066] | CI: [-0.2350, 0.7421] | CI: [-0.2690, 0.6889] |
| **State variability** | **state S** | *rho* = 0.37 | *rho* = 0.27 | *rho* = 0.09 | *rho* = -0.05 | *rho* = 0.19 | *rho* = 0.26 | *rho* = 0.23 | *rho* = -0.03 |
| | | *p* = 0.1213 | *p* = 0.2159 | *p* = 0.7048 | *p* = 0.8372 | *p* = 0.4182 | *p* = 0.2283 | *p* = 0.3774 | *p* = 0.8762 |
| | | CI: [-0.0645, 0.6990] | CI: [-0.2551, 0.7214] | CI: [-0.3707, 0.5427] | CI: [-0.4725, 0.3697] | CI: [-0.2557, 0.5799] | CI: [-0.2181, 0.6322] | CI: [-0.3094, 0.6789] | CI: [-0.4704, 0.3671] |
| | **state I** | *rho* = 0.39 | *rho* = 0.20 | *rho* = 0.37 | *rho* = -0.21 | *rho* = 0.33 | *rho* = 0.10 | *rho* = 0.36 | *rho* = -0.05 |
| | | *p* = 0.0637 | *p* = 0.3499 | *p* = 0.0860 | *p* = 0.3428 | *p* = 0.1162 | *p* = 0.6495 | *p* = 0.0888 | *p* = 0.8301 |
| | | CI: [0.0473, 0.6574] | CI: [-0.2173, 0.5519] | CI: [0.0210, 0.6464] | CI: [-0.6812, 0.2838] | CI: [-0.0488, 0.6632] | CI: [-0.3871, 0.4802] | CI: [-0.0562, 0.6859] | CI: [-0.5450, 0.5015] |

**Table S8.** Spearman's correlations between DFC parameters for not-centered data and 442 ROIs. Bold: significant correlation.

| | | Mean dwell time | | Prevalence S | Inter-transition interval | State variability | |
|---|---|---|---|---|---|---|---|
| **16-min data** | | **State S** | **State I** | **State S** | | **State S** | **State I** |
| **Mean dwell time** | **State S** | - | - | - | - | - | - |
| | **State I** | **rho = -0.4649 p = 0.0449** | - | - | - | - | - |
| **Prevalence** | **State S** | **rho = 0.655 p = 0.0023** | **rho = -0.7305 p = 0.0004** | - | - | - | - |
| **Inter-transition interval** | | **rho = 0.4965 p = 0.0306** | *rho* = 0.4246 *p* = 0.0700 | *rho* = -0.1115 *p* = 0.6495 | - | - | - |
| **State variability** | **State S** | *rho* = 0.3825 *p* = 0.1061 | **rho = -0.5719 p = 0.0105** | **rho = 0.4838 p = 0.0359** | *rho* = -0.2491 *p* = 0.3037 | - | - |
| | **State I** | **rho = -0.5772 p = 0.0097** | *rho* = 0.4351 *p* = 0.0626 | **rho = -0.7357 p = 0.0003** | *rho* = -0.1825 *p* = 0.4547 | *rho* = 0.0175 *p* = 0.9432 | - |
| **8-min data** | | | | | | | |
| **Mean dwell time** | **State S** | - | - | - | - | - | - |
| | **State I** | *rho* = -0.3846 *p* = 0.1755 | - | - | - | - | - |
| **Prevalence** | **State S** | *rho* = 0.4593 *p* = 0.1008 | **rho = -0.6088 p = 0.0237** | - | - | - | - |
| **Inter-transition interval** | | **rho = 0.7495 p = 0.0030** | *rho* = 0.1341 *p* = 0.6485 | *rho* = 0.1341 *p* = 0.6485 | - | - | - |
| **State variability** | **State S** | *rho* = 0.0725 *p* = 0.8083 | *rho* = -0.4066 *p* = 0.1505 | ***rho* = 0.7275 *p* = 0.0045** | *rho* = -0.3011 *p* = 0.2950 | - | - |
| | **State I** | *rho* = -0.4549 *p* = 0.1044 | *rho* = 0.3275 *p* = 0.2530 | ***rho* = -0.7363 *p* = 0.0038** | *rho* = -0.5165 *p* = 0.0616 | *rho* = -0.2308 *p* = 0.4265 | - |

**Table S9.** Spearman's correlations between DFC parameters for centered data and 442 ROIs. Bold: significant correlation.

| | | Mean dwell time | | Prevalence S | Inter-transition interval | State variability | |
|---|---|---|---|---|---|---|---|
| **16-min data** | | **State S** | **State I** | **State S** | | **State S** | **State I** |
| **Mean dwell time** | **State S** | - | - | - | - | - | - |
| | **State I** | rho = 0.41<br>p = 0.0501 | - | - | - | - | - |
| **Prevalence** | **State S** | rho = 0.19<br>p = 0.3758 | rho = -0.30<br>p = 0.1702 | - | - | - | - |
| **Inter-transition interval** | | **rho = 0.89**<br>**p = 0.0000** | **rho = 0.74**<br>**p = 0.0000** | rho = 0.09<br>p = 0.6716 | - | - | - |
| **State variability** | **State S** | rho = -0.08<br>p = 0.7032 | **rho = 0.59**<br>**p = 0.0030** | rho = -0.39<br>p = 0.0685 | rho = 0.18<br>p = 0.4090 | - | - |
| | **State I** | rho = 0.01<br>p = 0.9714 | rho = 0.34<br>p = 0.1092 | rho = -0.12<br>p = 0.5868 | rho = 0.16<br>p = 0.4601 | **rho = 0.58**<br>**p = 0.0038** | - |
| **8-min data** | | | | | | | |
| **Mean dwell time** | **State S** | - | - | - | - | - | - |
| | **State I** | **rho = 0.44**<br>**p = 0.0441** | - | - | - | - | - |
| **Prevalence** | **State S** | **rho = 0.69**<br>**p = 0.0005** | rho = 0.30<br>p = 0.1748 | - | - | - | - |
| **Inter-transition interval** | | **rho = 0.88**<br>**p = 0.0000** | **rho = 0.76**<br>**p = 0.0001** | **rho = 0.60**<br>**p = 0.0036** | - | - | - |
| **State variability** | **State S** | rho = -0.02<br>p = 0.9318 | rho = 0.21<br>p = 0.3399 | rho = -0.08<br>p = 0.7280 | rho = 0.11<br>p = 0.6103 | - | - |
| | **State I** | rho = 0.16<br>p = 0.4853 | rho = -0.10<br>p = 0.6426 | rho = 0.25<br>p = 0.2670 | rho = 0.02<br>p = 0.9478 | **rho = 0.50**<br>**p = 0.0204** | - |

Although we chose k=2 clusters out of theoretical considerations, we still evaluated whether our choice was optimal in terms of the silhouette statistic in session A (Fig. S1). Although the silhouette statistic results from k = 2 to 20 showed more fluctuations for the not-centered data than for the centered data in all the strategies, the results consistently showed an optimal cluster number of k = 2.

**Fig. S1.** Silhouette statistics for different time-length, atlases and not-centered/ centered data.

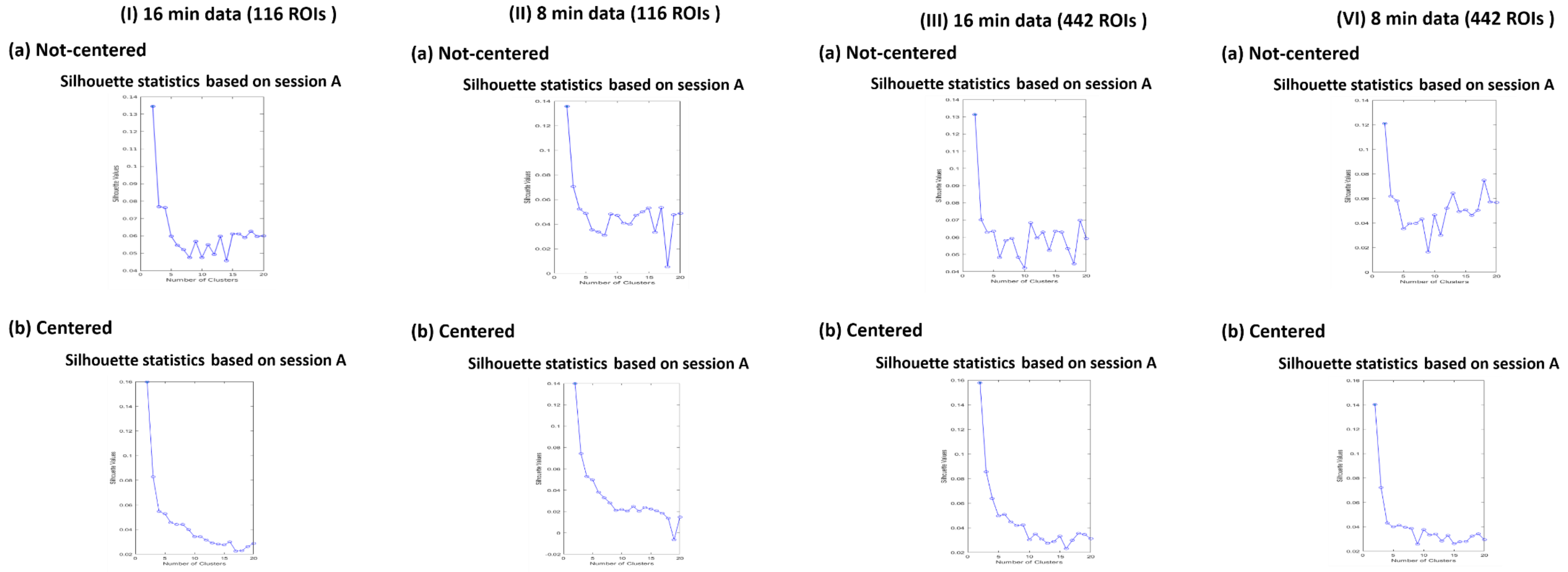

**Fig. S2.** Cluster centroids (k = 2) of the DFC matrices based on 442 ROIs for different time-length and not-centered/centered data. For session A, k-means clustering was used for the state assignment, while back projection results are shown for session B. Displayed are the group means of the within-run means (N1: visual network; N2: sensory-motor network; N3: dorsal attention network; N4: ventral attention network; N5: limbic network; N6: fronto-parietal network; N7: default mode network; N8: basal ganglia network; N9: cerebellum network).

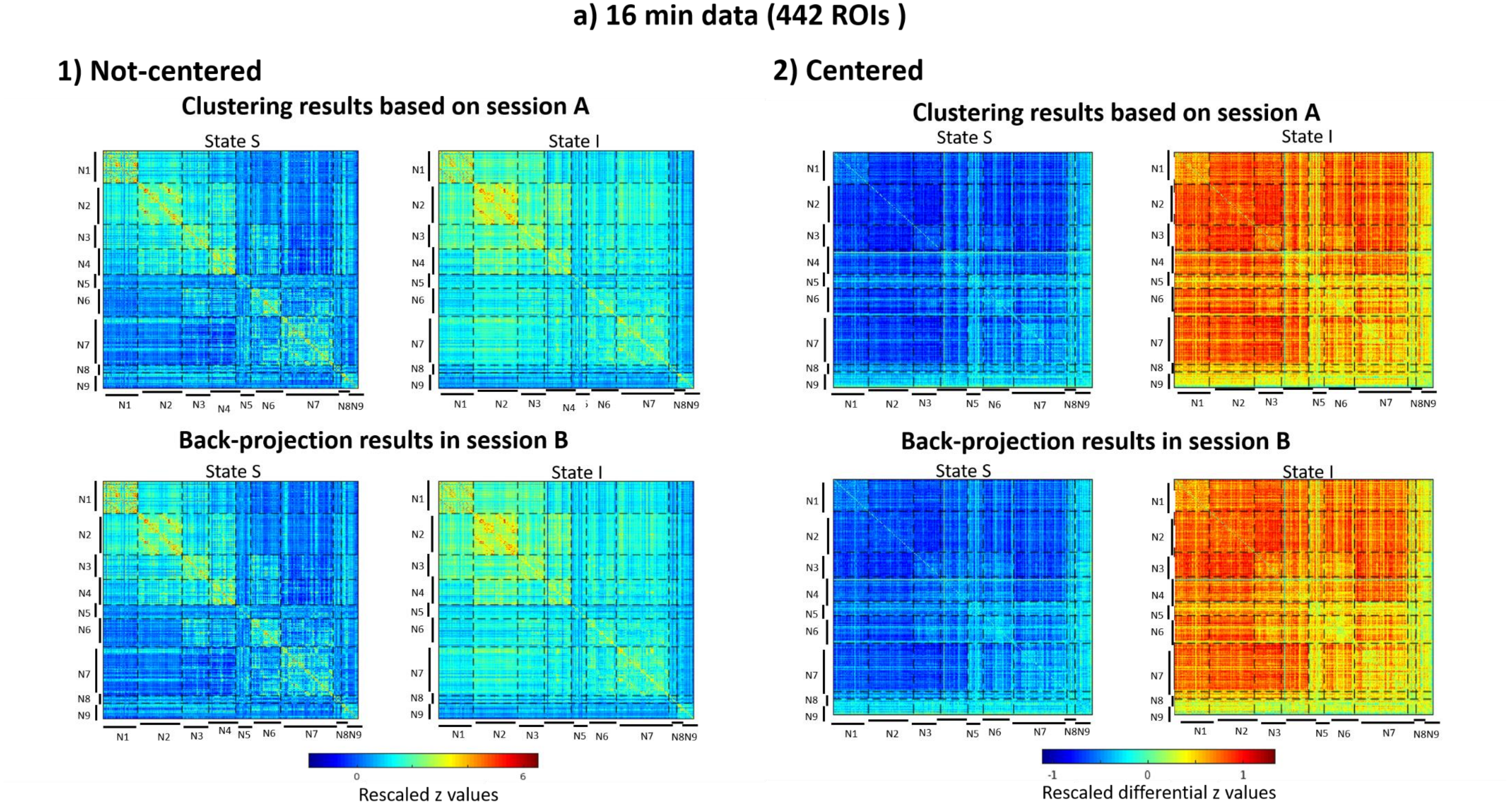

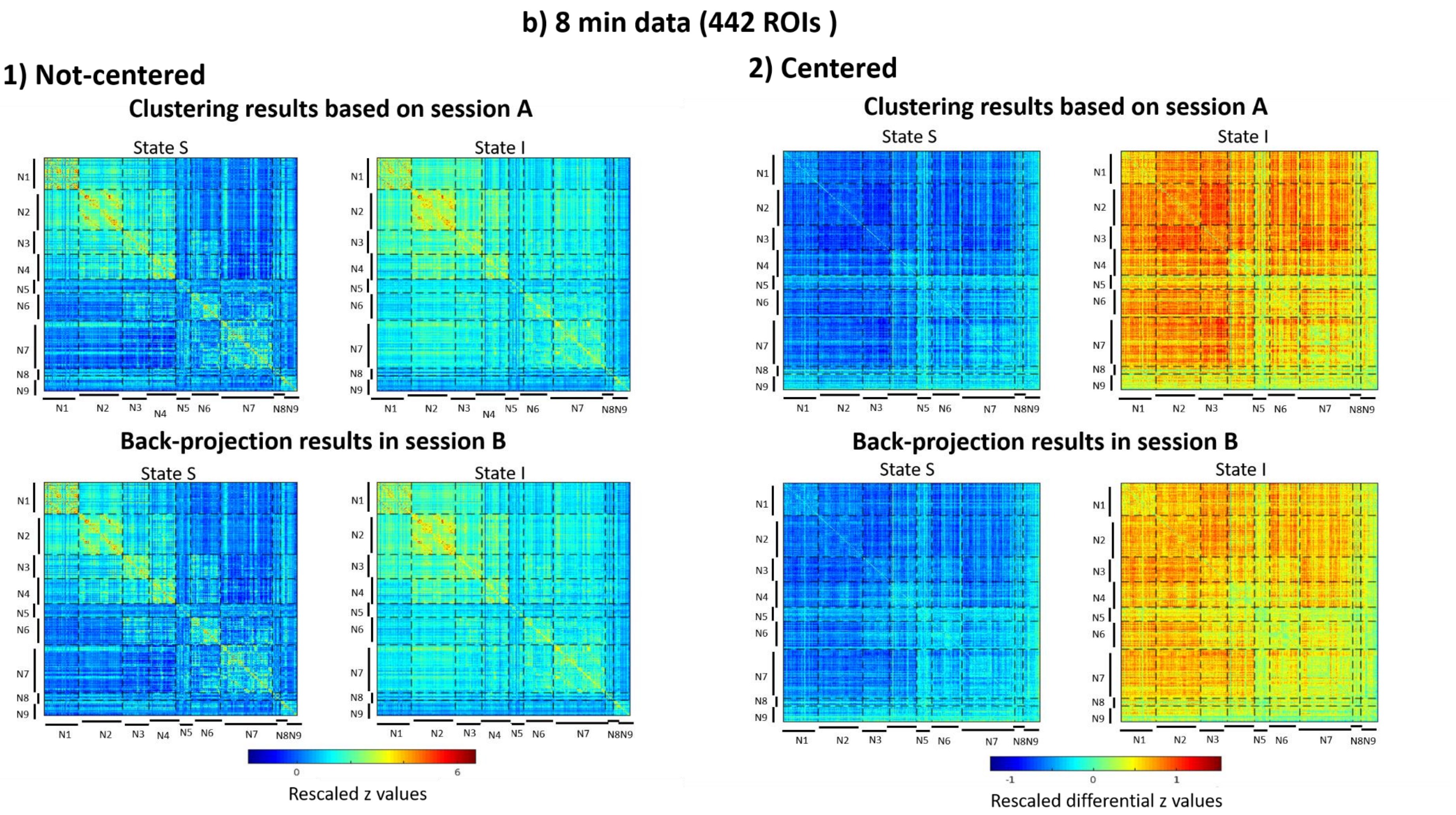

b) 8 min data (442 ROIs )
1) Not-centered
Clustering results based on session A
State S
State I
N1 N2 N3 N4 N5 N6 N7 N8 N9
Back-projection results in session B
State S
State I
0
6
Rescaled z values
2) Centered
Clustering results based on session A
State S
State I
Back-projection results in session B
State S
State I
-1
0
1
Rescaled differential z values

**Fig. S3.** Distance plots of all the frames to the two cluster centroids for different time-length, atlases and not-centered/centered data.

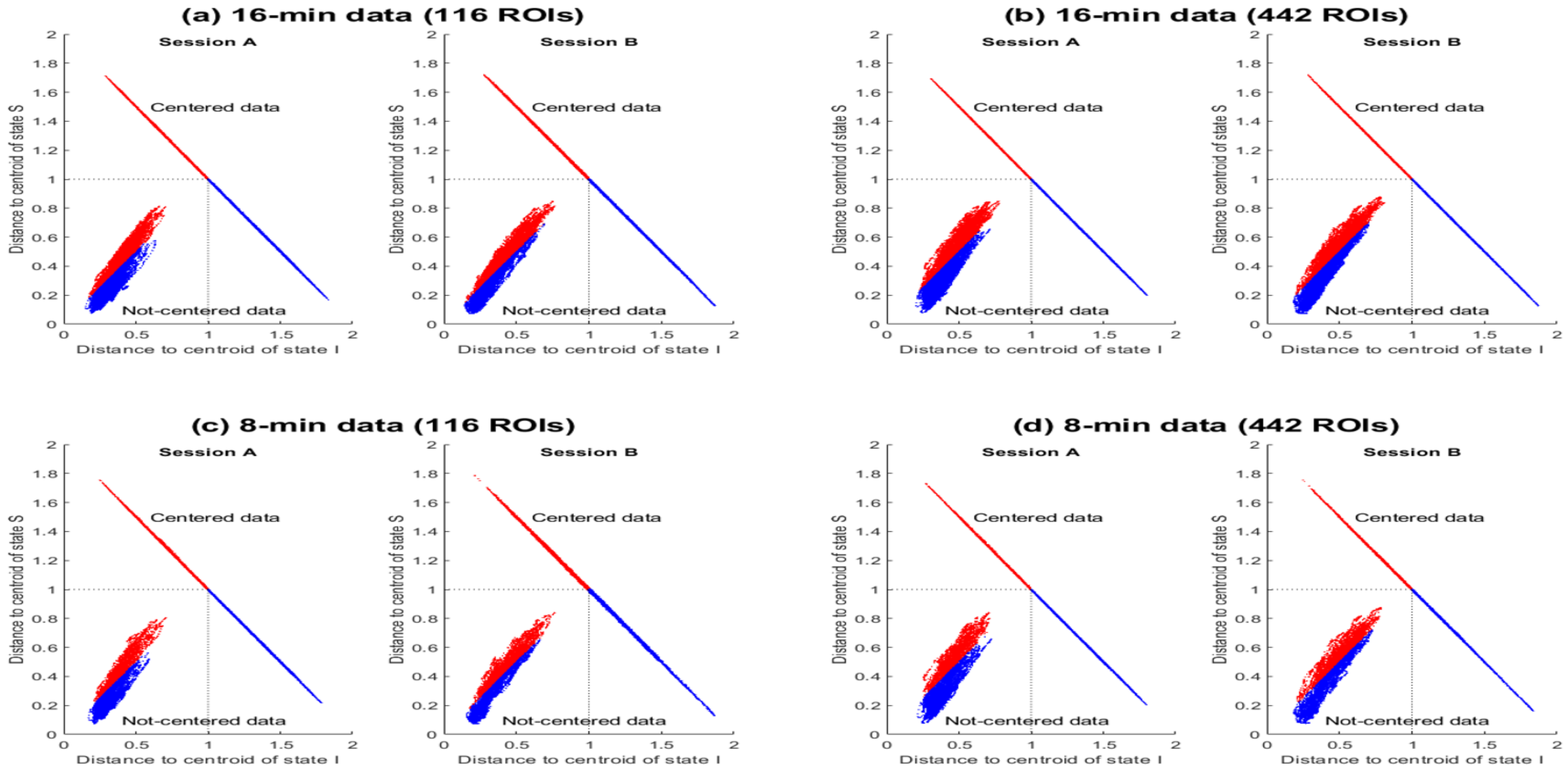

**Fig. S4.** Scatter plots of reliabilities of mean dwell time (*MDT*) and inter-transition time (*ITI*) in units of TR, and state variability (*Var*) based on different pipelines. Solid line: $p$ value < 0.05; dotted line: $p$ value > 0.05.

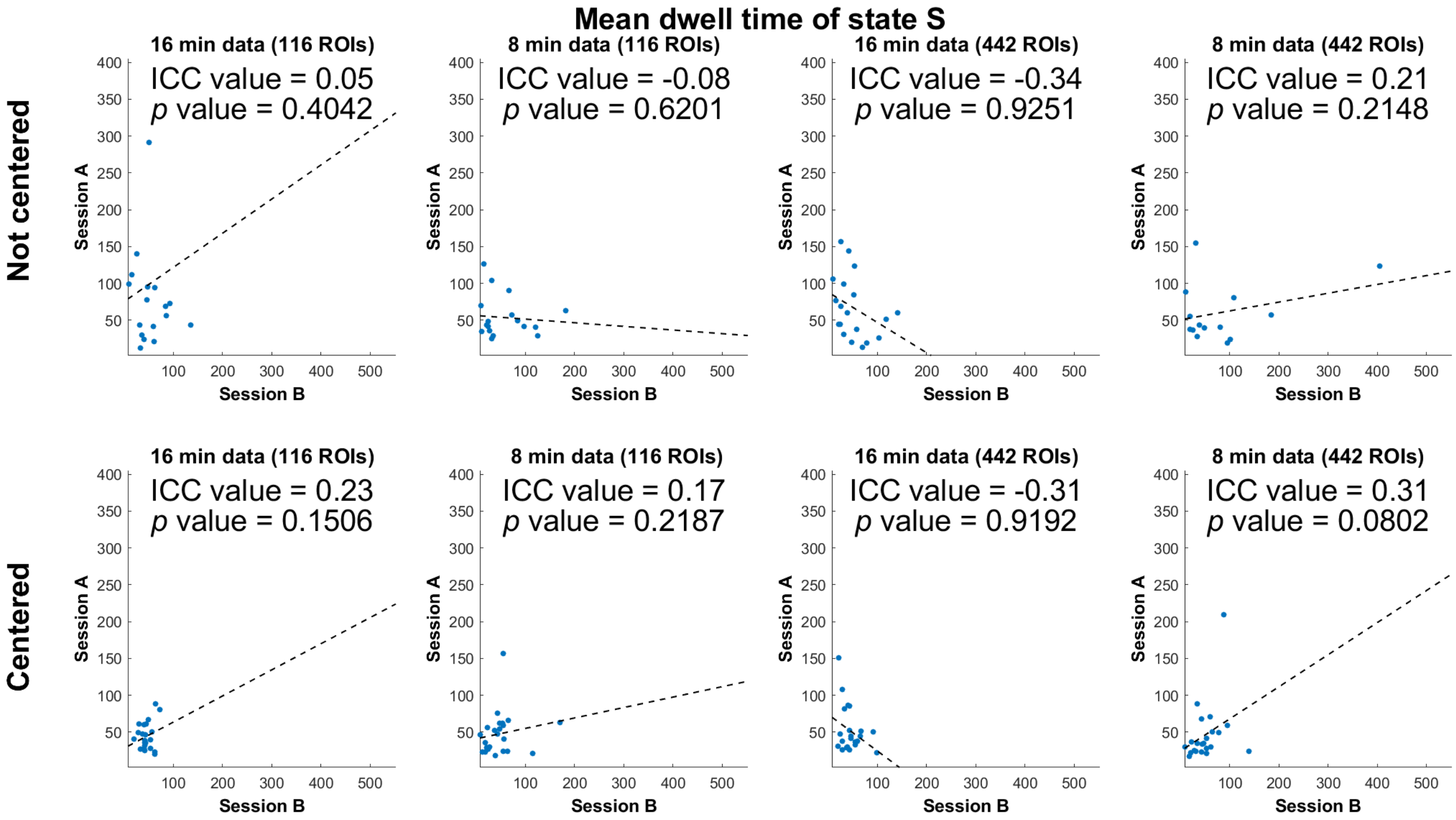

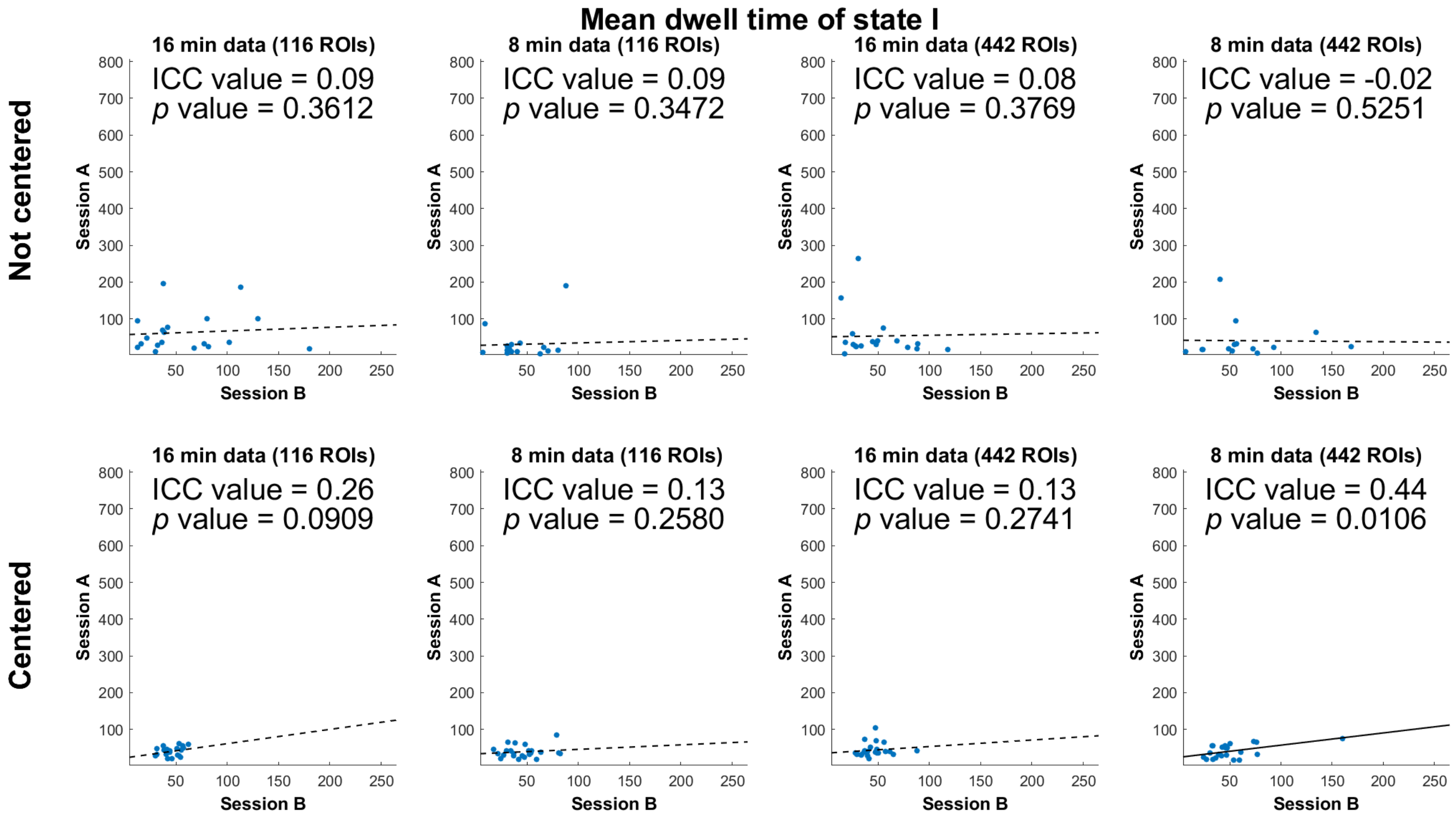
Mean dwell time of state I
Not centered
Centered
16 min data (116 ROIs)
8 min data (116 ROIs)
16 min data (442 ROIs)
8 min data (442 ROIs)
ICC value = 0.09
p value = 0.3612
ICC value = 0.09
p value = 0.3472
ICC value = 0.08
p value = 0.3769
ICC value = -0.02
p value = 0.5251
ICC value = 0.26
p value = 0.0909
ICC value = 0.13
p value = 0.2580
ICC value = 0.13
p value = 0.2741
ICC value = 0.44
p value = 0.0106
Session A
Session B
100
200
300
400
500
600
700
800
50
100
150
200
250

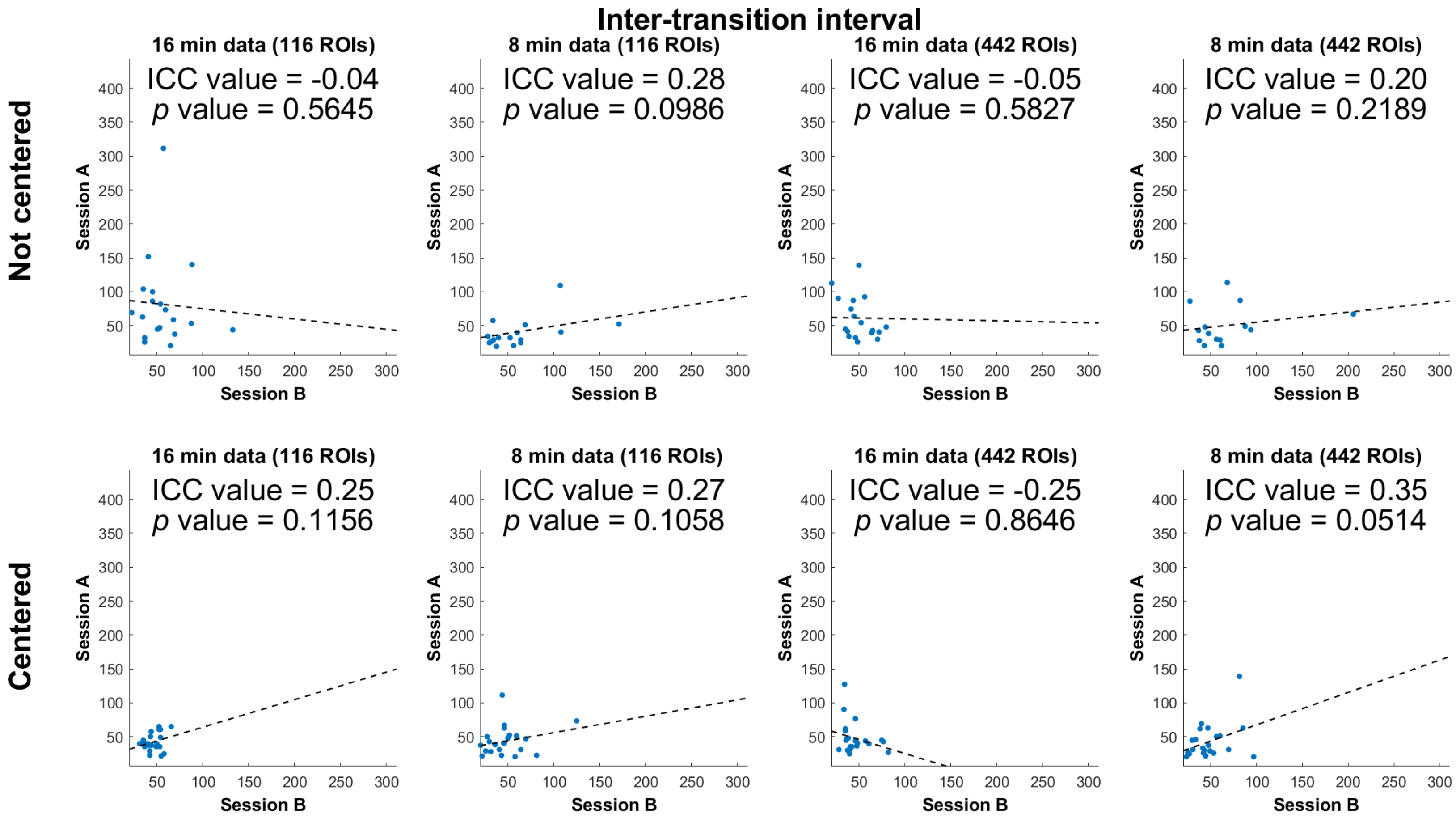
Inter-transition interval
Not centered
Centered
16 min data (116 ROIs)
ICC value = -0.04
p value = 0.5645
8 min data (116 ROIs)
ICC value = 0.28
p value = 0.0986
16 min data (442 ROIs)
ICC value = -0.05
p value = 0.5827
8 min data (442 ROIs)
ICC value = 0.20
p value = 0.2189
16 min data (116 ROIs)
ICC value = 0.25
p value = 0.1156
8 min data (116 ROIs)
ICC value = 0.27
p value = 0.1058
16 min data (442 ROIs)
ICC value = -0.25
p value = 0.8646
8 min data (442 ROIs)
ICC value = 0.35
p value = 0.0514
Session A
Session B

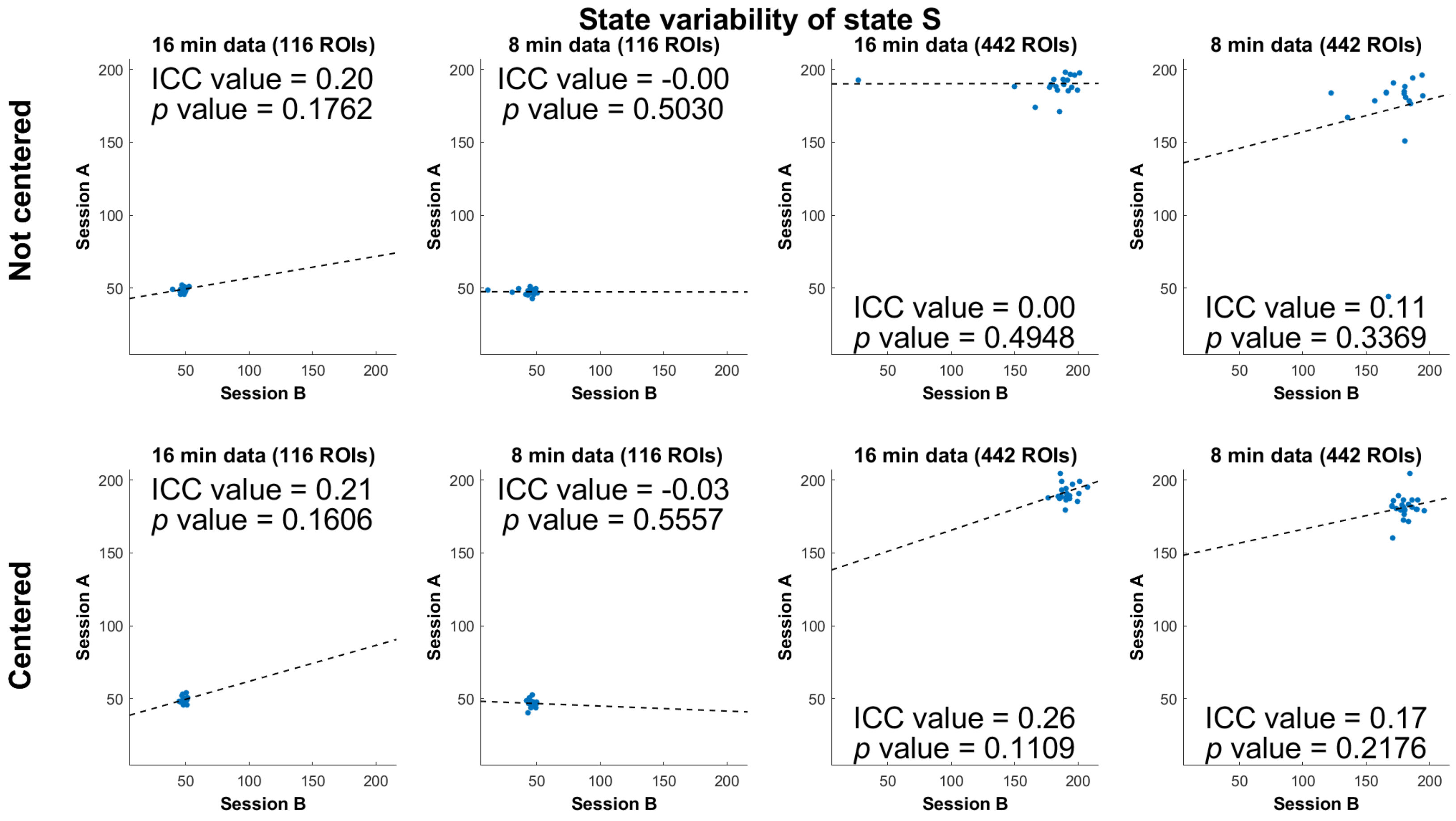
State variability of state S
Not centered
Centered
16 min data (116 ROIs)
ICC value = 0.20
p value = 0.1762
8 min data (116 ROIs)
ICC value = -0.00
p value = 0.5030
16 min data (442 ROIs)
ICC value = 0.00
p value = 0.4948
8 min data (442 ROIs)
ICC value = 0.11
p value = 0.3369
16 min data (116 ROIs)
ICC value = 0.21
p value = 0.1606
8 min data (116 ROIs)
ICC value = -0.03
p value = 0.5557
16 min data (442 ROIs)
ICC value = 0.26
p value = 0.1109
8 min data (442 ROIs)
ICC value = 0.17
p value = 0.2176
Session A
Session B

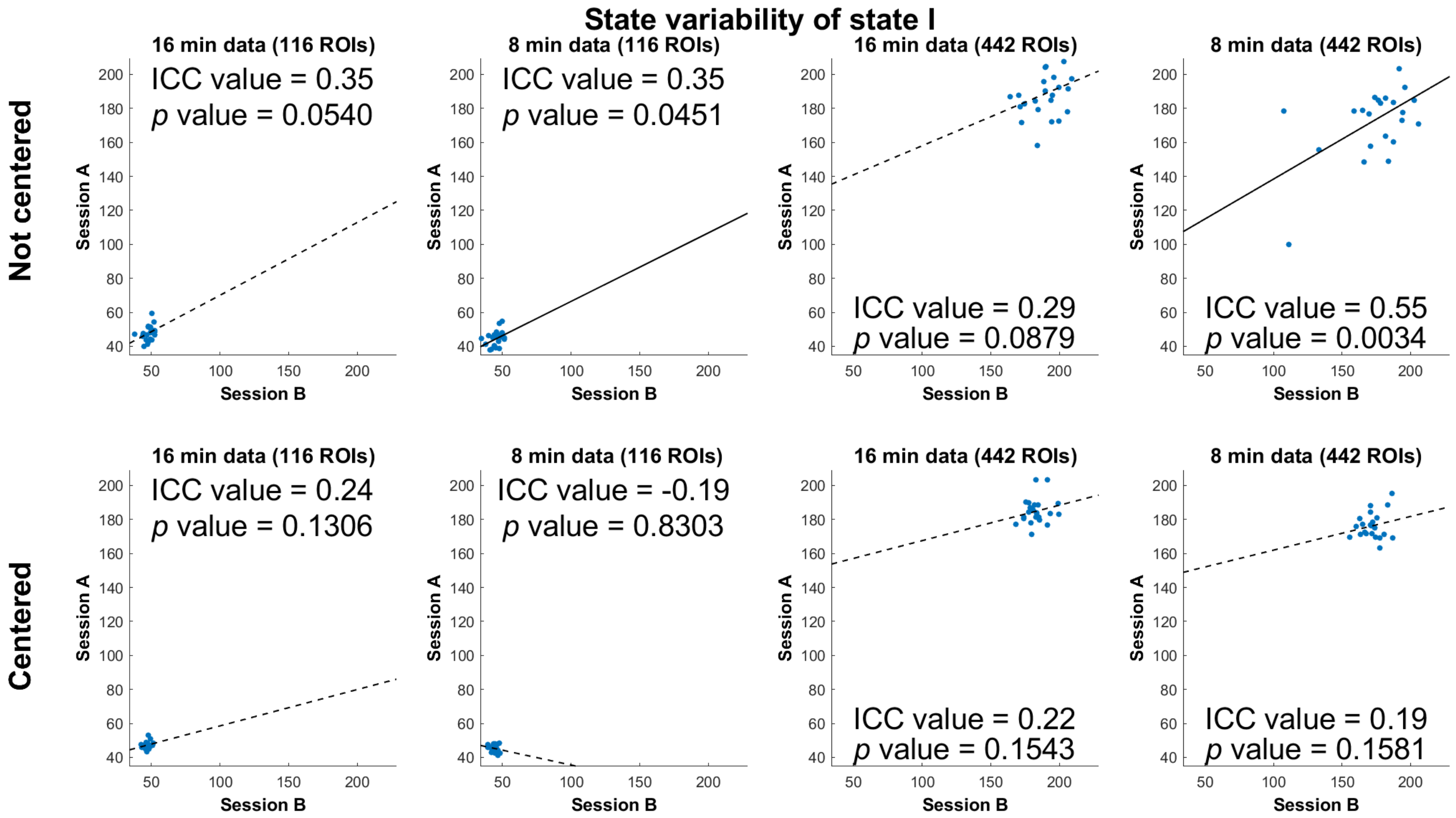
State variability of state I
Not centered
Centered
16 min data (116 ROIs)
ICC value = 0.35
p value = 0.0540
8 min data (116 ROIs)
ICC value = 0.35
p value = 0.0451
16 min data (442 ROIs)
ICC value = 0.29
p value = 0.0879
8 min data (442 ROIs)
ICC value = 0.55
p value = 0.0034
16 min data (116 ROIs)
ICC value = 0.24
p value = 0.1306
8 min data (116 ROIs)
ICC value = -0.19
p value = 0.8303
16 min data (442 ROIs)
ICC value = 0.22
p value = 0.1543
8 min data (442 ROIs)
ICC value = 0.19
p value = 0.1581
Session A
Session B

**Fig. S5.** Associations between sessions of fluctuations of all ROI-ROI connections (i.e., z-values) in the clusters after removing the session-mean matrices based on 8-min., not-centered data with 442 ROIs.

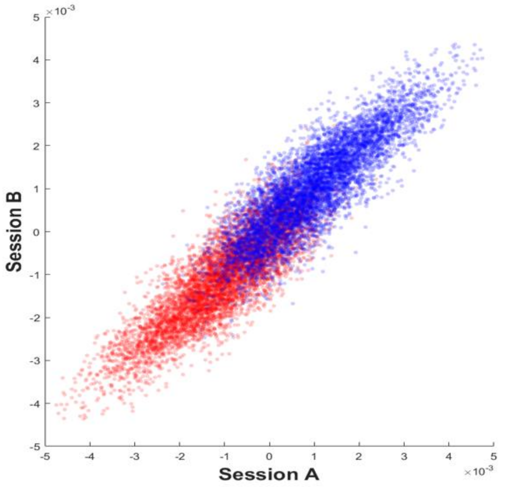


**1) Plots for the same ROI-ROI connectivity from State S (red) versus state I (blue) across the two sessions**

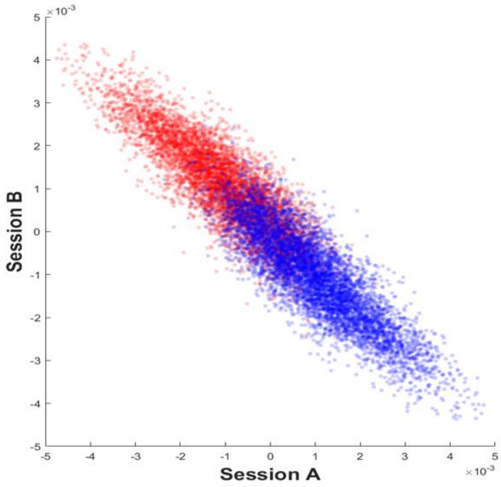


**2) Plots for the same ROI-ROI connectivity from unpaired states and sessions. Red: connectivity in state S in session A and states I in session B; blue: connectivity in state I in session A and state S in session B.**

**Fig. S6.** Plot of the distances of state matrices to the centroid of both states (16-min, 116 ROIs, session A) showing DFC example matrices (centered data) near the boundary between clusters and far away from the boundary. (state S – red; state I – blue)

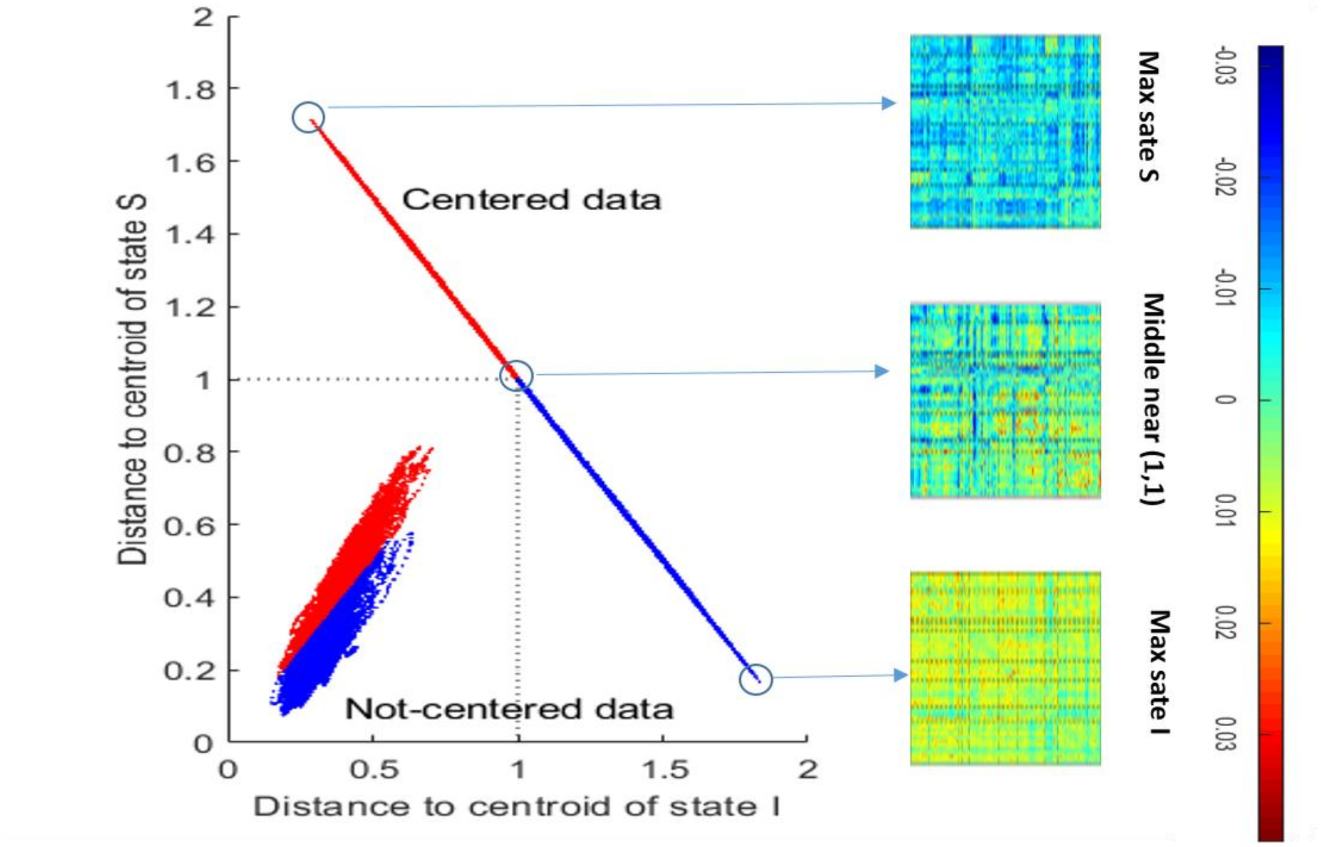
